# Palmitoylation of the Hemagglutinin of Influenza B virus by ER-localized DHHC enzymes 1, 2, 4 and 6 is essential for virus replication

**DOI:** 10.1101/2023.06.15.545201

**Authors:** Xiaorong Meng, Michael Veit

## Abstract

Covalent attachment of the fatty acids palmitate or stearate to the cytoplasmic domain of viral glycoproteins is often crucial for viral replication. This has previously been studied for the hemagglutinin (HA) of Influenza A virus and the responsible enzymes have been identified, but similar studies have not been performed with HA of Influenza B virus, which contains palmitate linked to two cysteines. We show here that the modification is essential for virus replication since exchange of both cysteines or the cysteine located at the end of the cytoplasmic tail prevented the generation of viable viruses. Viruses with an exchange of the membrane-distal cysteine rapidly reverted back to wild-type virus. Blocking exit of proteins from the endoplasmic reticulum (ER) revealed that palmitoylation of HA of Influenza B virus occurs in the ER, whereas acylation of HA of Influenza A virus also in the Golgi. Infecting cells deficient in DHHC palmitoyltransferases revealed that HA of Influenza B virus is acylated by the ER-localized DHHCs 1, 2, 4 and 6, which are thus different from the enzymes previously identified for acylation of HA of Influenza A virus. A comparison of predicted and experimentally determined protein structures suggests that the exclusive acylation of the HA of Influenza B virus with palmitate is not a function of the responsible DHHCs and that the transmembrane region may be critical for the acylation of HA of Influenza A and B virus by different DHHCs.

**Importance:** Influenza viruses are a public health concern since they cause seasonal outbreaks and occasionally pandemics. Our study investigates the importance of a protein modification called "palmitoylation" in the replication of Influenza B virus. Palmitoylation involves attaching fatty acids to the viral protein hemagglutinin, and has previously been studied for Influenza A virus. We found that this modification is essential for the Influenza B virus to replicate, as mutating the sites where palmitate is attached prevented the virus from generating viable particles. Our experiments also showed that this modification occurs in the endoplasmic reticulum. We identified the specific enzymes responsible for this modification, which are different from those involved in palmitoylation of HA of Influenza A virus. Overall, our research illuminates the similarities and differences in fatty acid attachment to HA of Influenza A and B virus and identifies the responsible enzymes, which might be promising targets for antiviral therapy.

## Introduction

Protein palmitoylation is a post-translational modification that involves the addition of a fatty acid, mainly palmitate or stearate, to specific cysteine residues in a membrane protein. This modification is critical for a variety of cellular processes, including protein trafficking, signaling, and membrane association [1]. Palmitoylation is also observed in many enveloped viruses, including Influenza viruses, where it might be crucial for the replication and spread of the virus [2]. The enzymes responsible for palmitoylation are the DHHC proteins, a family of polytopic membrane proteins containing an Asp-His-His-Cys (DHHC) motif as the catalytic center embedded within a cysteine-rich domain (CRD) in one of their cytoplasmic domains. DHHCs exhibit a two-step reaction mechanism, a fatty acid is first transferred from the lipid donor acyl-CoA to the cysteine of the DHHC motif (autoacylation) and subsequently to a cysteine of the substrate protein. Humans have 23 different DHHC proteins, which have different substrate specificities that only partially overlap. Besides the CRD, there is little sequence conservation between DHHC proteins, and they usually possess four, only some have more transmembrane regions [3, 4]. The majority of DHHC proteins localize to the Golgi, a small number remains in the endoplasmic reticulum (ER) or is targeted to the plasma membrane, but cell type specific differences have been described [5, 6].

The crystal structures of two DHHC-proteins, human DHHC 20 and zebrafish DHHC 15 have been determined. These structures reveal that the four transmembrane helices form a tent-like structure, with the DHHC motif located at the membrane-cytosol interface. The autoacylated form of DHHC 20 contains the irreversible inhibitor bromo-palmitate attached to Cys 156 of the DHHC motif. The fatty acid is inserted into a hydrophobic cavity formed by all four transmembrane regions. At the narrow end of the cavity Ser 29 forms a hydrogen bond with Tyr 181, which effectively closes the groove. Ser 29 and Tyr 181 are not conserved between DHHC proteins; all DHHCs contain either two bulky residues, one bulky and one small or two small amino acids at the homologues position. It was hypothesized that the presence of certain amino acids (large or small) determines the lipid binding specificity of a DHHC enzyme [7].

Influenza viruses are a significant public health concern, causing seasonal outbreaks and occasional pandemics. Influenza A viruses have the ability to infect a wide range of animal species, including pigs and horses, but wild birds are considered their natural reservoir. Influenza B and C viruses, on the other hand, are primarily human pathogens, with the latter usually causing only mild symptoms [8, 9].

Hemagglutinin (HA) is the glycoprotein responsible for binding to sialic acid-containing receptors on host cells and for fusion of viral and cellular membranes, allowing the virus to enter and infect the cell, in both Influenza A and B viruses. Influenza C virus, on the other hand, has a different glycoprotein called hemagglutinin-esterase-fusion (HEF) protein, which also has receptor-destroying activity in addition to its role in membrane fusion. HA and HEF are trimeric type I transmembrane glycoproteins with an N-terminal signal peptide, a long extracellular domain containing the functional activities, a transmembrane domain, and a short cytoplasmic tail [10, 11]. The proteolytic cleavage of HA into two subunits, HA1 and HA2, is a crucial step for Influenza virus replication. The cleavage allows the virus to expose the fusion peptide in the HA2 subunit, which is necessary for the virus to enter the host cell. The proteolytic cleavage of HA can occur either intracellularly, in the trans-Golgi network by cellular Furin-proteases, or at the plasma membrane, by proteases of the TMPRSS family [12].

HA and HEF are acylated, but with a different number and type of fatty acid. Influenza B virus HA contains palmitate at two cytoplasmic cysteines, whereas HEF of Influenza C virus contains stearate attached to one cysteine located at the cytosolic site of the transmembrane region. HAs of Influenza A virus having one transmembrane and two cytoplasmic cysteines contain both palmitate and stearate, but the latter is exclusively attached to the cysteine positioned in the TMR of HA [13, 14]. Acylation of Flu A HA occurs on the trimeric HA, but before it acquires Endo-H resistant carbohydrates, suggesting that it occurs in the ‘late’ ER, in the ER–Golgi intermediate compartment or in cis-Golgi cisternae [15].

The role of acylation of Influenza A HA has been studied extensively, revealing its essential function in virus replication. Palmitoylation of the two cytoplasmic cysteines is crucial, while stearoylation of the third cysteine has only a moderate effect [16-18]. Acylation facilitates the enrichment of the protein in small nanodomains of the plasma membrane, affecting both the assembly of virus particles and the membrane fusion activity of HA [19-24]. Notably, the acylation sites are conserved across all HA subtypes and variants, despite the high variability of the molecule [25]. In contrast, the removal of the stearoylated cysteine from HEF of Influenza C virus has only modest effects on virus growth and membrane fusion activity [26]. Although the essential role of HA acylation has been demonstrated in Influenza A virus, no studies with recombinant viruses have investigated its function in Influenza B virus. Interestingly, expression of HA lacking the C-terminal acylation site can promote hemifusion, but not pore formation. However, deleting both acylation sites or the cytoplasmic tail has no effect on membrane fusion, and it remains unknown whether these small effects measured in an artificial system affect virus replication [27, 28].

While the acylation of many glycoproteins from different virus families is known to be crucial for virus replication, only a few of the DHHC enzymes responsible for this modification have been identified. The spike protein of SARS-CoV-2 is predominantly acylated by DHHC 20, with a contribution by DHHC 9 and other DHHCs [29-31]. In our previous work, we identified DHHC 2, 8, 15 and 20 as enzymes responsible for acylation of HA of Flu A, but we found that the same set of enzymes had little or no effect on acylation of HA of Flu B and HEF of Flu C [32]. In this study, we investigate whether the acylated cysteines are essential for Influenza B virus replication and aimed to identify the enzymes responsible for palmitoylating HA. These enzymes might be putative drug targets, since their inhibition could mitigate virus spread with potentially little effect on cell viability [33].

## Results

### Palmitoylation of HA is essential for replication of Influenza B virus

We replaced the two cysteines in the cytoplasmic tail of HA of the Influenza B virus reference strain Lee by serine, either separately (mutants Ac1 and Ac2) or together (Ac1+2, Fig. 1A). The mutant HA plasmid, together with seven additional plasmids encoding the other viral proteins, was transfected into a co-culture of 293T and MDCKII cells after the two nucleotide alterations TGT to AGC were confirmed by sequencing. Three days later the supernatant was removed and the remaining cells were subjected to immunofluorescence with antibodies against HA to show that transfection was successful (Fig. 1B, upper row). An aliquot of the supernatant (P1 virus) was amplified in fertilized eggs (P2 virus) and subsequently in MDCKII cells (P3). The cellular supernatant (undiluted for Ac2 and Ac1+2, 1:10 diluted for Ac1 and 1:100 diluted for wt-infected cells) was used to inoculate MDCKII cells. Immunofluorescence revealed that many cells were infected with Flu B wt, fewer cells were infected with Ac1 and no cells were infected with Ac2 and Ac1+2 (Fig. 1B, lower row). The second aliquot from the transfection was passaged three times in MDCKII cells and the resulting supernatants were subjected to HA assay. The wild-type virus from passage 3 (P3 virus) and 4 (P4 virus) produced HA titres of 2E7, while no agglutination was measurable with P4 virus from Ac1, Ac2 and Ac1+2 (Fig. 1C). We performed two more transfections with a subsequent amplification in eggs or in MDCK cells, but were never able to detect infectious virus particles for Ac2 and Ac1+2. Note also that Ac1 could only be successfully rescued if the supernatant of the transfection was subsequently amplified in embryonated eggs. To test for the genetic stability of the inserted mutation, Flu B Ac1 from embryonated eggs (P2 virus) was amplified two more times in MDCK II cells and after each amplification step, cDNA was prepared and ∼300 nucleotides at the 5’ end of the HA gene were sequenced to analyse whether the mutation is still present (Fig. 2A). Whereas the P2 virus still contains the codon AGC introduced into the plasmid, the sequencing chromatogram of the P3 virus reveals two peaks at position one (A or T) and two at position three (T or C) of this codon. It thus contains a mixture of viruses having either a Ser (AGC, AGT) or a Cys (TGC, TGT) in HA. After a further amplification in MDCK II cells the resulting P4 virus contains exclusively TGT which is identical to the codon present in HA wt (Fig. 2A). We conclude that a fatty acid attached to the membrane-near Cys is beneficial for virus growth, since viruses having this amino acid rapidly outgrow viruses with a Ser. Since a simultaneous exchange of two nucleotides in one amplification step is unlikely, the sequence of reversions from AGC (Ser) to AG**T** (Ser) and to **T**GC (Cys) in passage 3 and to TGT (Cys) in passage 4 seems most probable. The original codon TGT is apparently also preferred over the codon TGC, which also codes for cysteine.

**Fig 1.**
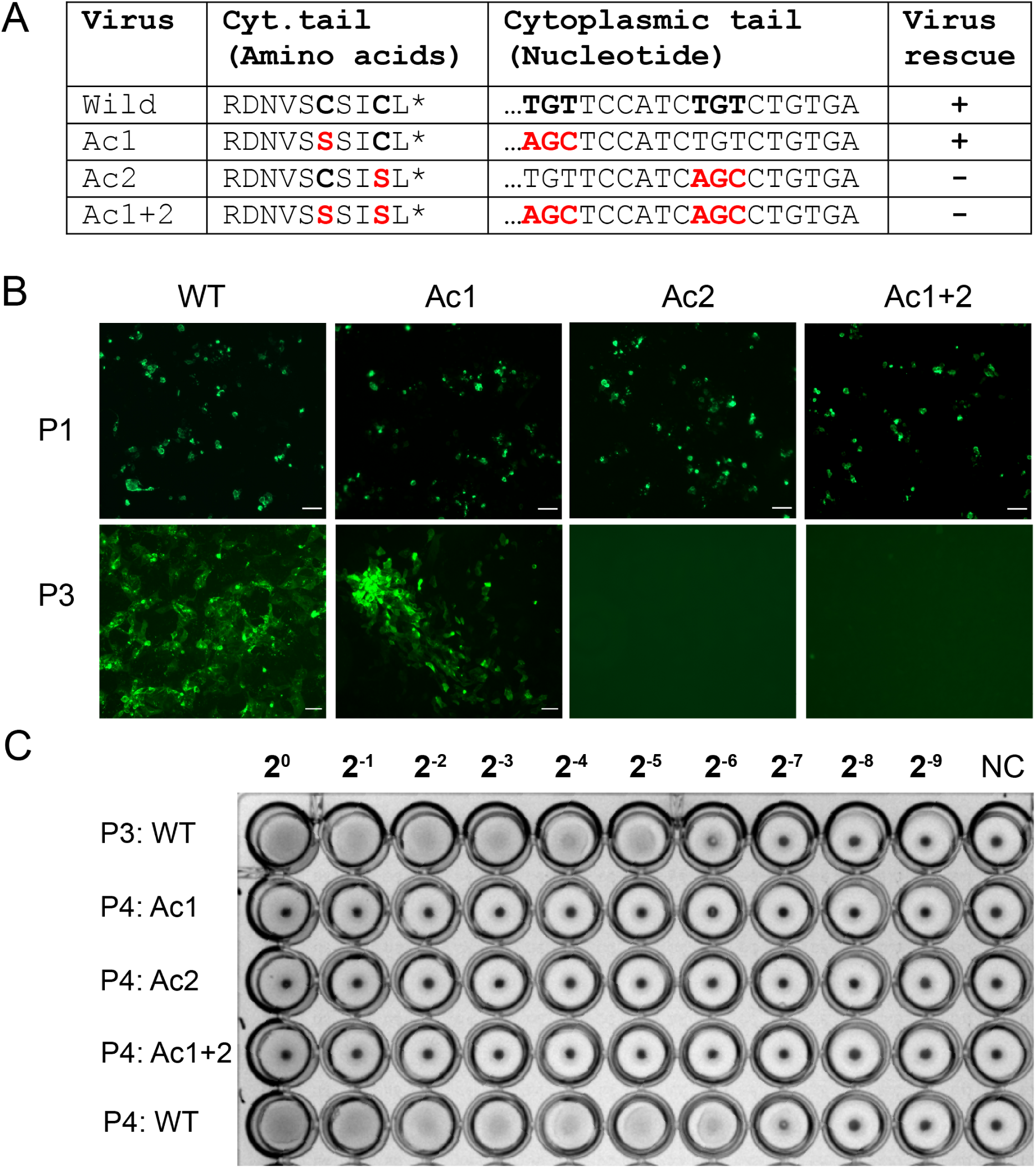
Palmitoylation of HA is essential for Influenza B virus replication. (A) Amino acid and nucleotide sequence of HA from the Influenza B virus strain B/Lee/40. Acylated cysteines and the encoding nucleotide triplets are in bold. The mutations are highlighted in red. An asterisk indicates the stop codon. + and - indicates whether or not virus could be rescued. (B) Upper row: Immunofluorescence of a co-culture of 293T and MDCKII cells transfected with seven Flu B plasmids and the indicated HA plasmid. Lower row: Immunofluorescence of MDCKII cells infected with P3 virus, which was generated by amplification of the transfection supernatants in eggs and subsequently in MDCKII cells. The scale bar is 100 µm. (C) HA-assay with P3 and P4 virus, which was generated by two and three amplification steps of the transfection supernatants only in MDCKII cells. Note, that we could rescue the Flu B Ac1 only, if the supernatant of the transfection was first amplified in embryonated eggs.

**Fig. 2.**
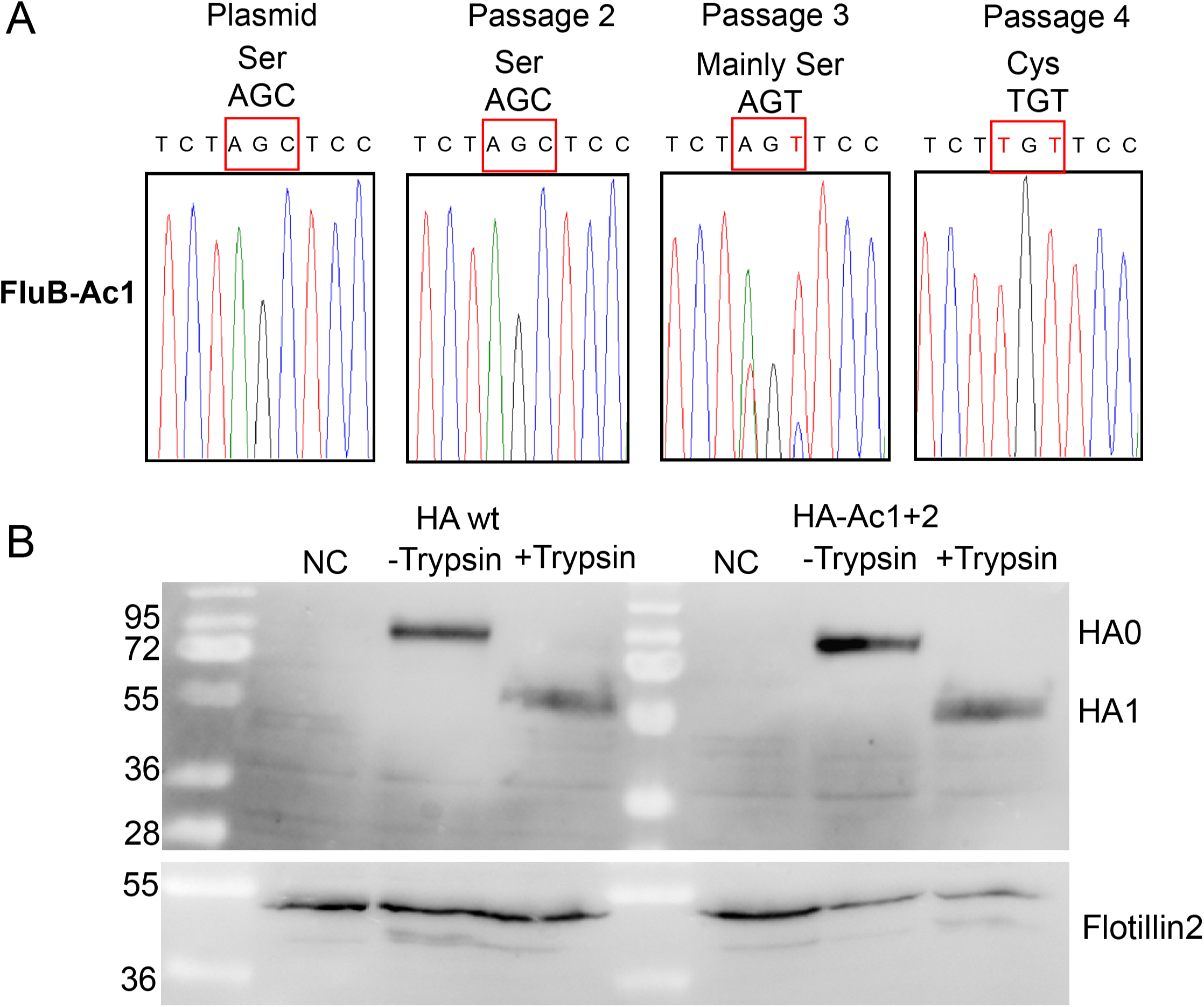
Flu B Ac1 rapidly reverted back to the wild-type sequence. (A) Sequence chromatograms of the plasmid used to rescue Ac1 and of the recombinant virus after one passage in embryonated eggs (P2 virus) and one (P3) or two (P4) passages in MDCKII cells. (B) Expression of HA wt and HA Ac1+2 and cell surface trypsinisation. HA wt and HA Ac1+2 were expressed in 293T cells, which were treated after 48h with trypsin for 30 minutes. Cells were lysed and subjected to western blot with anti-HA antibody. Antibodies against the cellular protein flotillin2 were used as loading control.

Palmitoylation may be crucial for the stability and intracellular trafficking of viral glycoproteins. [29]. In order to determine if the inability to rescue the Ac1+2 recombinant is due to impaired transport of HA to the budding site, we expressed HA Ac1+2 and HA wt in 293T cells, which were subsequently subjected to trypsin treatment. Only when HA is exposed at the cell surface does the enzyme cleave it into its subunits HA1 and HA2. In the absence of trypsin, a HA band of similar intensity is detected for HA wt and HA Ac1+2. In the presence of trypsin HA0 wt as well as HA0 Ac1+2, is completely cleaved into HA1 and HA2, the latter band is not visible since the monoclonal antibody binds to the HA1 subunit (Fig. 2B). Thus, palmitoylation does not affect the stability of HA or its intracellular transport, as previously reported for HA of Flu A and HEF of Flu C [26, 34].

### Palmitoylation of HA of Influenza B virus occurs in the endoplasmic reticulum

In an effort to reduce the number of DHHCs potentially implicated in palmitoylation of HA of Flu B, we investigated in which compartment along the exocytic the modification happens and whether it varies from the compartment where HA of Flu A is acylated. We employed Brefeldin A (BFA), a drug that prevents proteins from leaving the ER, to study this [35, 36]. Although no tool exists to test directly for protein exit from the ER, transport of HA to the medial Golgi can be monitored using endoglycosidase-H (Endo-H), which removes carbohydrates of the high mannose type. Posttranslational proteolytic cleavage of HA into HA1 and HA2 subunits is another criterion for vesicular transport. Despite having a monobasic cleavage site, HA of Flu B is partially proteolytically activated in MDCK II cells by a transmembrane serine protease [37-39]. Similarly, HA of the Influenza A strain WSN is also partially cleaved in these cells, either by endogenous proteases or facilitated by recruitment of serum plasminogen and its conversion to the protease plasmin [40-42].

First, we explored various experimental setups and discovered that a modest dose of BFA, given three hours after infection and present the entire time, prevented HA transport while also allowing enough HA to be synthesized to test it for acylation. In untreated cells Endo-H removes carbohydrates from approximately half of the uncleaved HA molecules but not from HA1 because cleavage of HA occurs after the acquisition of Endo-H resistant carbohydrates. Because the antibody only binds to the HA1 subunit, HA2 is not visible in the blot. Only a small part of HA molecules is cleaved in the presence of BFA, and both uncleaved HA and HA1 are entirely sensitive to Endo-H digestion, showing that cleavage occurs under BFA treatment before HA develops Endo-H resistant carbohydrates (Fig. 3A). The same experiment performed with Influenza A virus strain WSN provides almost identical results: BFA prevents cleavage of HA and also the conversion of uncleaved HA to an Endo-H resistant form (Fig. 3B). Also take note of the fact that uncleaved HA0 in BFA-treated cells of both Influenza A and B virus has a slightly lower molecular weight than uncleaved HA0 in untreated cells. This is because ER-located glycoproteins have shorter, trimmed carbohydrate chains, while in the Golgi the side chains are extended.

**Fig 3.**
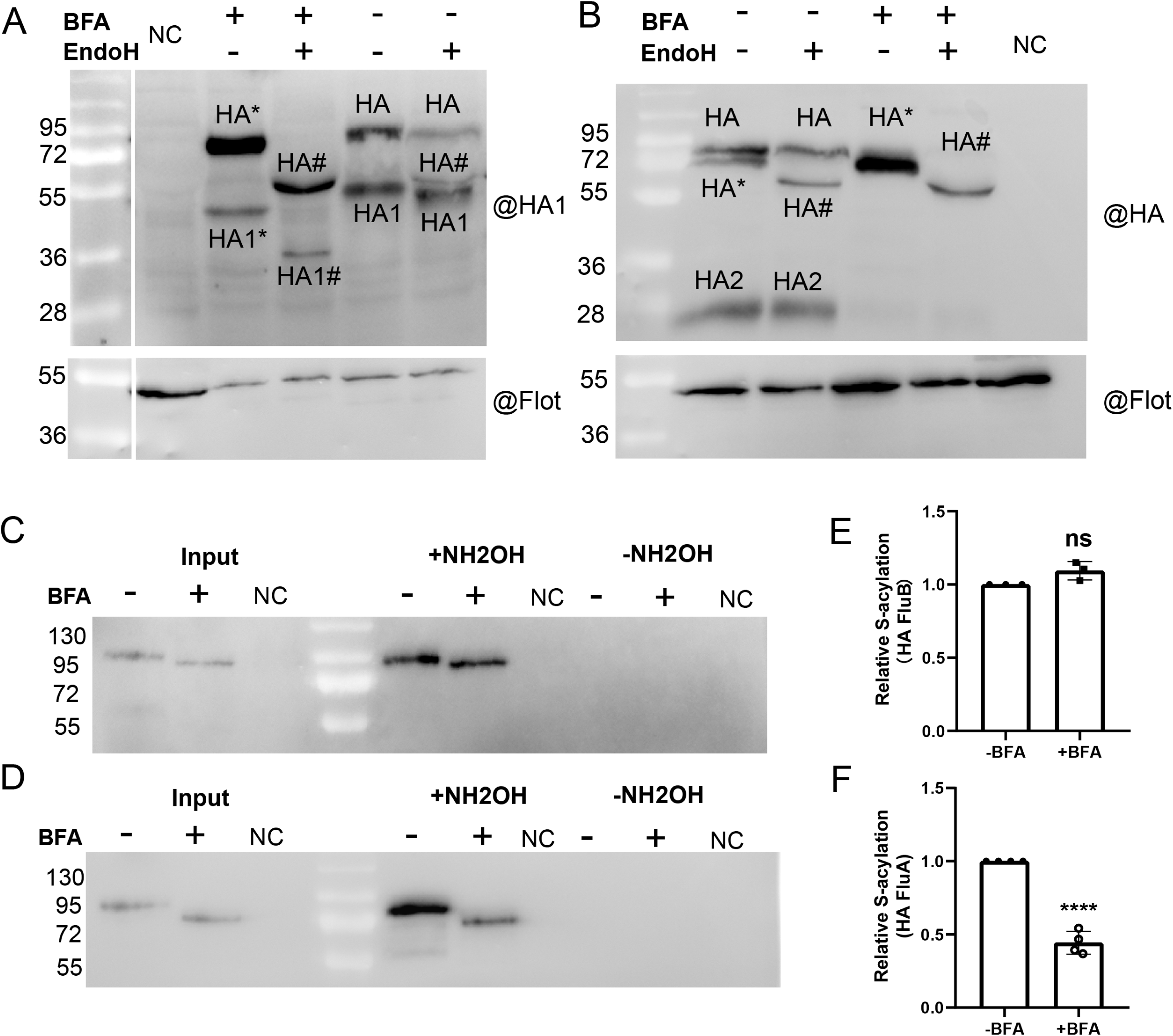
Palmitoylation of HA of Flu B occurs in the ER. (A,B) Validating the effect of brefeldin A. MDCKII cells were infected with Influenza B virus (A) or Influenza A virus (B) at an moi of 1. 3hours later BFA (3µg/ml) was added and cells were incubated for 24 (A) or 12 (B) hours. Cells were lysed using NP40 lysis buffer and treated or not treated with Endo H glycosidase. Lysates were then subjected to western blot with antibodies against Influenza B Virus HA (A) or Influenza A virus HA (B). Labelling of bands: HA: uncleaved HA, HA1, HA2: cleaved subunits, #: HA with carbohydrates removed by Endo-H digestion. *: Glycosylated HA with lower molecular weight due to the BFA-induced transport block. Note that HA bands in untreated cells are weaker since the protein was lost due to virus budding, which does not occur if HA transport is blocked. (C, D) S-acylation level of HA of Flu B (C) or Flu A (D) in the presence of BFA. MDCKII cells were infected with Influenza B virus (C) or Influenza A virus (D) and treated with BFA as described. Cell lysates were then subjected to Acyl-RAC assay, which is based on pull-down of proteins with thiol-active resins. Input: aliquot of the cell lysate to compare expression levels. NH_2_OH: samples treated (+) or not treated (-) with hydroxylamine to cleave thioester-bound fatty acids. Note that a larger amount of the cell lysate was used from all samples from non-treated cells to account for protein loss caused by virus budding. No HA1 of Flu B and very little HA2 of Flu A were detected in these blots. (E, F) Quantification of three independent experiments with Influenza B (E) and four with Influenza A (F) virus. Densities of the +NH_2_OH bands were divided by the densities of the input bands and normalized to experiments without BFA. Unpaired t-test was applied for statistical analysis, ns, not significant, ****, P < 0.0001 versus wild type without BFA

Once the experimental parameters for blocking HA transport had been established, we compared the acylation of HA in BFA-treated and untreated cells. We employed the Acyl-RAC (resin-assisted capture) assay, which makes use of thiol-reactive resins to capture SH-groups that had just been released as a result of hydroxylamine cleavage of thioester linkages. MDCKII cells were infected with the Influenza B virus in the presence or absence of BFA, lysed, and a 10% aliquot was used to compare the levels of HA expression (input). The remaining protein sample was equally split, one aliquot was treated with hydroxylamine (+NH_2_OH), the other aliquot was treated with Tris– HCl (-NH_2_OH) to assess the binding specificity of the resin. All samples were then subjected to western blotting with antibodies against HA (Fig 3C, D). Quantification of the results from three independent experiments revealed that acylation of HA of Flu B was undiminished, whereas acylation of HA of Flu B was reduced to ∼45% with very little variation between experiments (Fig, 3E, F). In these experiments, only a very small fraction of HA of Flu A was cleaved in untreated cells and HA of Flu B revealed no cleavage (not shown). We have no conclusive explanation for the inter-experimental variability other than the MDCK-II cell line being a mixture of three different cell types that differ in the expression of proteolytic enzymes [39]. We conclude that acylation of HA of Flu A Influenza HA begins in the ER but continues in the Golgi, which is consistent with previous findings [15]. In contrast, acylation of Flu B HA occurs exclusively in the ER.

### HA of Influenza B virus is acylated by DHHC enzymes 1, 2, 4 and 6

To identify the DHHCs involved in acylation of HA of Influenza B virus we used commercial HAP1 cells, where various ZDHHCs were knocked-out using the CRISPR/Cas9 technology. Wild-type HAP1 cells express all human ZDHHCs, except ZDHHC 19 [32]. They can be infected with Influenza B virus and synthesize HA, which is transported to the plasma membrane as demonstrated by the binding of erythrocytes to infected cells. However, HAP1 cells do not release virus particles into the medium as analysed by hemagglutination and plaque assays, as already observed for Influenza A virus (not shown). Note also that cleavage of HA was not observed in any experiment with HAP1 cells.

In the first screening experiments we used cell lines where the ER-located DHHCs 1, 2, 4 and 6, the Golgi-located DHHCs 5, 7, 8, 15, 16 have been inactivated [5, 6]. The HAP1 cells were infected with virus, lysed 24 hours later and subjected to Acyl-RAC to determine whether acylation of HA is reduced in any of the infected cells. The results of two independent experiments are shown in Fig. 4A and B and their quantification (densities of the +NH_2_OH bands relative to densities of the input bands) in Fig. 4C. In both experiments acylation of HA was quite markedly reduced only in ΔDHHC 2 and ΔDHHC 4 cells and slightly in ΔDHHC 1 cells. Most other knock-out cell lines revealed a large variation between both experiments, except for ΔDHHC 8 cells, which exhibit a small increase.

**Fig. 4.**
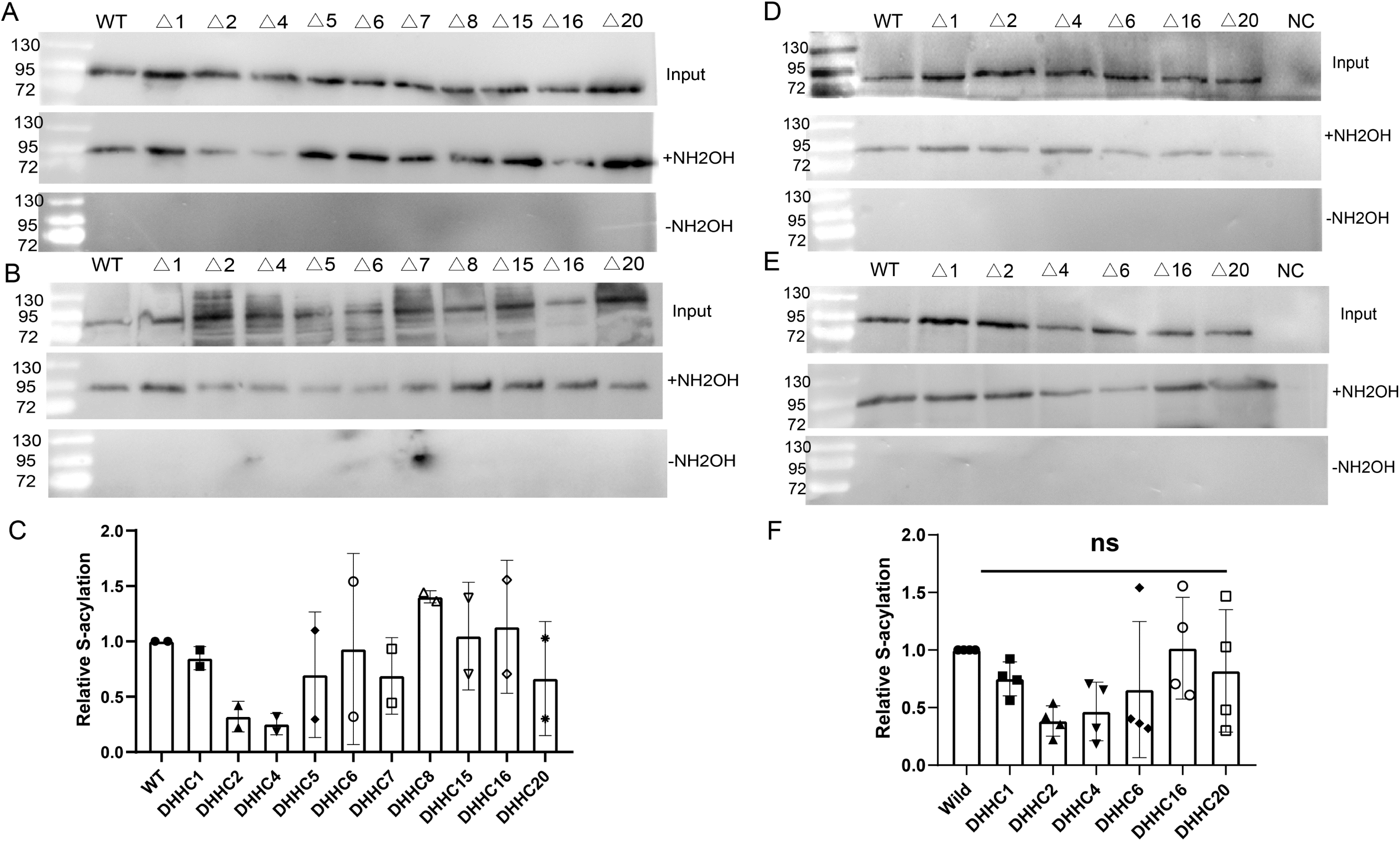
Testing HA acylation in cells deficient in ER- and Golgi-located DHHCs. (A, B, D, E) HAP1 cells where the indicated DHHC genes were knocked-out using the CRISPR/Cas9 technology were infected with Influenza B virus at an MOI of 1. 24 hours later, acylation of HA was analysed using Acyl-RAC. Input: aliquot of the cell lysate to compare expression levels. NH_2_OH: samples treated (+) or not treated (-) with hydroxylamine to cleave thioester-bound fatty acids. (C) Quantification of data from A and B. (D) Quantification of data from A, B, D and E. The mean ± SD is shown. One-way ANOVA followed by multiple comparison Dunnett test was applied for statistical analysis. ns, not significant compared to WT.

In the next screening round, we used the ΔDHHC 2 and ΔDHHC 4 cell lines, the ΔDHHC 1 and ΔDHHC 6 cells, where the other ER-located DHHCs have been inactivated and ΔDHHC 16 and ΔDHHC 20 cells. The results of two independent experiments show again a marked reduction in acylation of HA in ΔDHHC 2 and ΔDHHC 4 cells and a small reduction in ΔDHHC 1 cells. HA acylation was also markedly reduced in both experiments with ΔDHHC 6 cells, but no consistent effect was seen in ΔDHHC 16 and ΔDHHC 20 cells (Fig. 4D-F). However, quantification of the four experiments performed so far showed no statistically significant differences compared to wild-type cells for any of the knockout cells.

To further statistically validate the results, we performed the same experiment two more times with ΔDHHC 1, ΔDHHC 2, ΔDHHC 4 and ΔDHHC 6 cells. To exclude that proteins were lost during sample preparation, we used antibodies against the cellular palmitoylated protein flotillin 2 as an internal control (Fig. 5A and B). Acylation of HA is reduced in all four knockout cells, most obviously in ΔDHHC 2 and ΔDHHC 4 cells and quantification of all six experiments showed a statistically significant difference for these cells (Fig. 5C). In five of the six experiments, infection of ΔDHHC 6 cells showed a substantial decrease in acylation of HA. However, since the first experiment showed an increase in acylation, the evaluation of all experiments revealed no statistically significant difference to wt cells. When the first experiment is removed from the analysis, the average reduction in acylation is now significant and at the same level as in the ΔDHHC 2 and ΔDHHC 4 cells, namely a reduction to about 40 %. Similarly, the small reduction (10 %) in acylation of HA in ΔDHHC 1 cells is now also statistically significant (Fig. 5D). We conclude that DHHC 1, 2, 4 and 6 are involved in acylation of HA of Flu B.

**Fig. 5.**
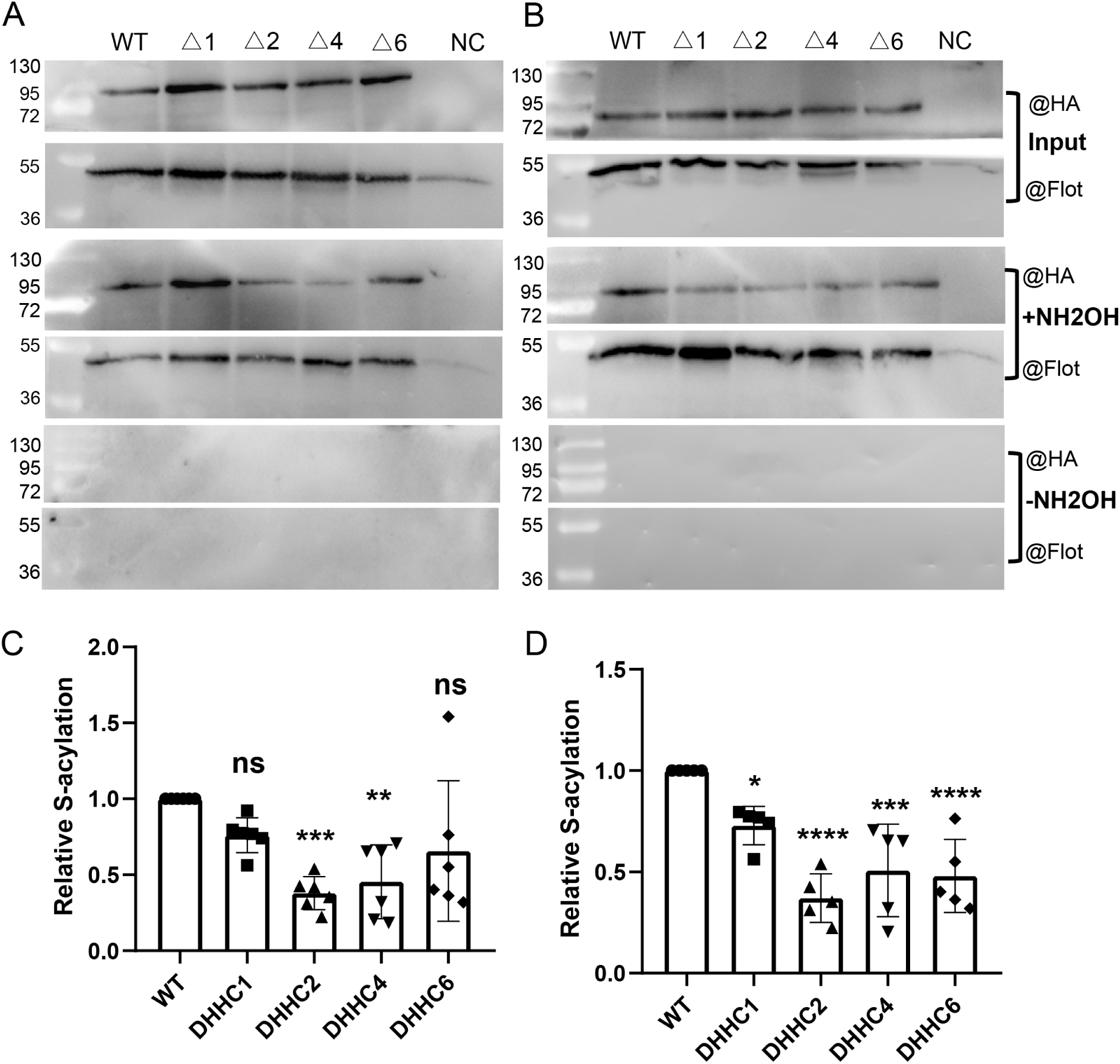
HA of Influenza B virus is acylated by DHHC 1, 2, 4 and 6. (A, B) HAP1 cells where the indicated DHHC genes were knocked-out using the CRISPR/Cas9 technology were infected with Influenza B virus at an MOI of 1. 24 hours later, acylation of HA was analysed using Acyl-RAC. Input: aliquot of the cell lysate to compare expression levels. NH_2_OH: samples treated (+) or not treated (-) with hydroxylamine to cleave thioester-bound fatty acids. (C) Quantification of data from all six experiments in Fig. 3 and 4. The mean ± SD is shown. One-way ANOVA followed by multiple comparison Dunnett test was applied for statistical analysis. ns, not significant, **, P <0.01, ***, P <0.001 versus WT. (D) Quantification of data from five experiments, excluding data from Fig. 3A. The mean ± SD is shown. One-way ANOVA followed by multiple comparison Dunnett test was applied for statistical analysis. *, P <0.05, ***, P <0.001, ****, P <0.0001 versus WT.

### DHHC 1, 2, 4 and 6 are located in the ER in cultured cells and expressed in epithelia cells of the human respiratory tract

DHHC 4 and 6 are typical ER-localized transmembrane proteins with a C-terminal lysine-based retention signal and DHHC1 remains in the ER by an unknown mechanism [43, 44]. DHHC 2 associates with the plasma membrane, recycling endosomes, and vesicular structures in neurons [45] and in T-cells it localizes primarily to the endoplasmic reticulum and Golgi apparatus [46]. We investigated the intracellular localization of DHHC 1, 2, 4 and 6 by confocal microscopy in transfected BHK cells, which were subsequently stained with an ER-marker (S1 Fig.). Quantification revealed the highest Pearsońs correlation coefficient (∼70) for DHHC 1, 4 and 6, but the value for DHHC 2 is only slightly lower (∼65). In contrast, DHHC 3, which resides mainly in the cis-Golgi [5] revealed a different, perinuclear staining pattern and a lower Pearsońs correlation coefficient (∼45). In addition, HA of Flu B shows substantial overlap with DHHC 1, 2, 4 and 6 in double transfected BHK cells (S2 Fig.). We also investigated localisation of DHHC 2, 4 and 6 in A549 cells, a model for alveolar Type II pulmonary epithelium, where Influenza viruses replicate (Fig. 6). We calculated the same Pearson correlation coefficient as in BHK cells, i.e. ∼70 for DHHC 4 and 6 and only slightly lower (∼65) for DHHC 2. We conclude that the DHHCs responsible for the acylation of HA of Influenza B virus are located in the ER, which is consistent with the intracellular site of its acylation.

**Fig. 6.**
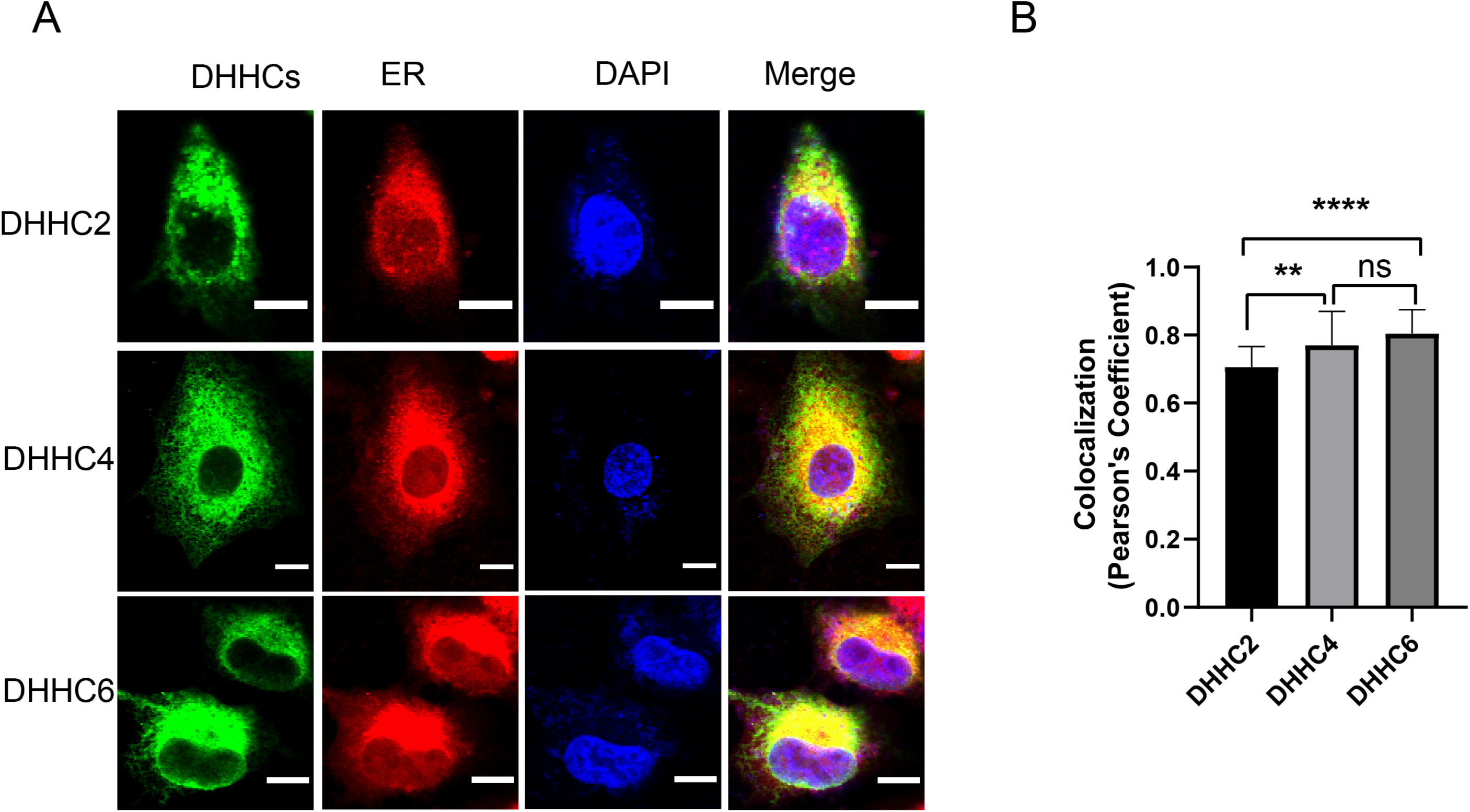
Co-localisation of DHHC 2, 4 and 6 with an ER-marker in human lung A549 cells. (A) Confocal microscopy of A549 cells transfected with the indicated mouse DHHCs fused to a HA tag. Cells were permeabilized and stained with antibodies against the HA-tag (green), with the ER staining kit (red), and with DAPI (blue) to highlight the nucleus. The scale bar is 10 µm. (B) Quantification of co-localization from at least 40 cells with Pearson’s correlation coefficient method using the JACoP plugin of the Fiji software. One-way ANOVA followed by multiple comparison Tukey test was applied for statistical analysis. ****, P <0.0001 DHHC 2 versus DHHC 6; **, P <0.01 2 versus DHHC 4. DHHC 4 and 6 has no significant differences from each other.

Influenza viruses replicate in epithelia cells of the upper and lower respiratory tract, as well as in alveolar cells of the lung in more severe cases. It is therefore interesting to determine whether these cells express the DHHCs involved in HA acylation. We used "CellPalmSeq," a curated RNAseq database of palmitoylating enzyme expression in human tissues and cells, for this purpose. [47]. The expression of DHHC 4 and 6 in the whole lung tissue is much higher than that of DHHC 1 and 2. The enzyme with the highest expression in single cells of the lung is DHHC 4, followed by DHHC 6. Except for alveolar type 1 and 2 cells, where DHHC 2 is most abundant, DHHC 2 is the third most expressed DHHC of the four. DHHC 1 is expressed at the lowest level and not at all in alveolar cells (S1 table). We hypothesise that DHHC 4 and 6 are responsible for the majority of acylation reactions of Flu B HA in human lung cells. It should be highlighted, nevertheless, that the stability and activity of the enzyme are just as important as the degree of RNA expression.

### The shape of HÁs transmembrane region might explain the acylation by different DHHCs

The HAs of Influenza A and B viruses are acylated by different DHHCs, only DHHC 2 has an effect on both proteins. We were therefore interested in whether the transmembrane regions or the cytoplasmic domains of the two HA proteins have peculiar structural features that could act as recognition motifs for their respective DHHC enzymes. Apart from the cryo-EM structure of a group 1 HA, in which, however, only the outer part of the transmembrane region was resolved, no other experimentally determined structures are available for full-length HA proteins [48]. Similarly, the alphafold2 database does not contain predicted HA structures [49]. We therefore used a colab version of alphafold2 multimer to predict the structures of the transmembrane region and cytoplasmic tail of HA of Flu B and of one group 1 and one group 2 HA of Flu A [50]. To estimate the accuracy of the prediction, alphafold2 supplies a per residue confidence metric called the predicted local distance difference test (pLDDT), which indicates the confidence in the local structure prediction. The pLDDT scores are displayed on the predicted structure in rainbow colours, ranging from red (high confidence) to blue (low confidence). The cytoplasmic tails have a very low pLDDT score below 50, which might suggest that they are intrinsically disordered. The transmembrane region, especially the inner part is predicted with a higher confidence score of 60 to 70 suggesting that at least the structural elements are predicted correctly (S3 Fig, left part). This is consistent with CD spectroscopy of purified peptides, that also revealed a trimeric helix for the transmembrane region of HA [51]. A surface representation of the transmembrane region allows to locate the amino acids exposed at the moleculés surface which might function as binding site of DHHC enzymes (S3 Fig, right part). However, the TMRs of HA of Flu B and Flu A revealed a similar pattern: large hydrophobic amino acid (Leu, Ile) are followed by two or three amino acids with -OH groups (Ser, Thr), which are followed by five or six hydrophobic amino acids, one or two of which are aromatic residues (Phe, Tyr, Trp). Looking only at the protein backbone, we see that the three helices in the HA of Flu B are straight and parallel to each other, whereas the helices of the HA s of Flu A are slightly bent. The kink in the helix is caused by a Gly localised in the lower part of the TMR. In the Cryo-EM structure of HA the region following the Gly splays out from the upper part of the transmembrane region [48]. Since a glycine is not present in the TMR of HAs from Flu B, its kink-inducing capacity is a feature that distinguishes HA from Flu A and Flu B.

### Exclusive attachment of palmitate to the HA of Flu B virus is not encoded in the responsible DHHCs

The crystal structure of the autoacylated form of DHHC 20 provided a model for the fatty acid specificity of the acylating enzymes [7]. The hydrogen bond established between Ser29 in TM1 and Tyr181 in TM3 in DHHC 20 closes the narrow end of the cavity in which the fatty acid is inserted. It was proposed that the specificity of a DHHC enzyme’s fatty acid binding depends on whether large or small amino acids are present at this position, which we investigated for DHHC 2, 4 and 6. Crystal structures are not available for any of these DHHCs, but the alphafold2 database contains predicted models with a per-residue confidence score (pLDDT) of >90 for the transmembrane regions, which are thus equivalent to experimentally determined structures. Other parts of the enzymes, especially domains down- or upstream of the four transmembrane regions, have a pLDDT score below 50 and hence are likely to be intrinsically disordered (Fig. 7, left part). The predicted models were superimposed with the structure of DHHC 20 to identify the crucial amino acids (Fig. 7, middle part). All three DHHCs, especially DHHC2 align very well with DHHC 20, especially over the entire transmembrane domain, including the cysteine-rich domain. DHHC 2 also possess a Ser in TM1 and a Tyr in TM3, which can be perfectly superimposed on their DHHC 20 counterparts (Fig. 7, right part). In addition, DHHC 2 contains a Tyr (instead of a Phe) in TM2, which forms an additional hydrophilic interaction with Ser in TM1 sealing the cavity further. DHHC 4 also contains a Ser/Tyr pair at the end of the hydrophobic cavity, but the order is reversed compared to DHHC 20. Tyr 80 is in TM1 and Ser204 in TM3 and both residues form a hydrophilic interaction. DHHC 6 contains Cys34 instead of Ser29 in TM1 and Gly154 instead of the bulky Tyr in TM3 and this pair of small amino acids is highly unlikely to prevent access of stearate into the hydrophobic cavity. The helix is only tightly sealed further up by a layer of hydrophobic amino acids. We conclude that the specific attachment of palmitate to HA of Flu B is not encoded in the DHHCs required for its acylation. Longer fatty acids, such as stearate, can likely be accommodated by DHHC 2, 4 and 6, which can transfer them to the HA of Influenza B virus.

**Fig 7.**
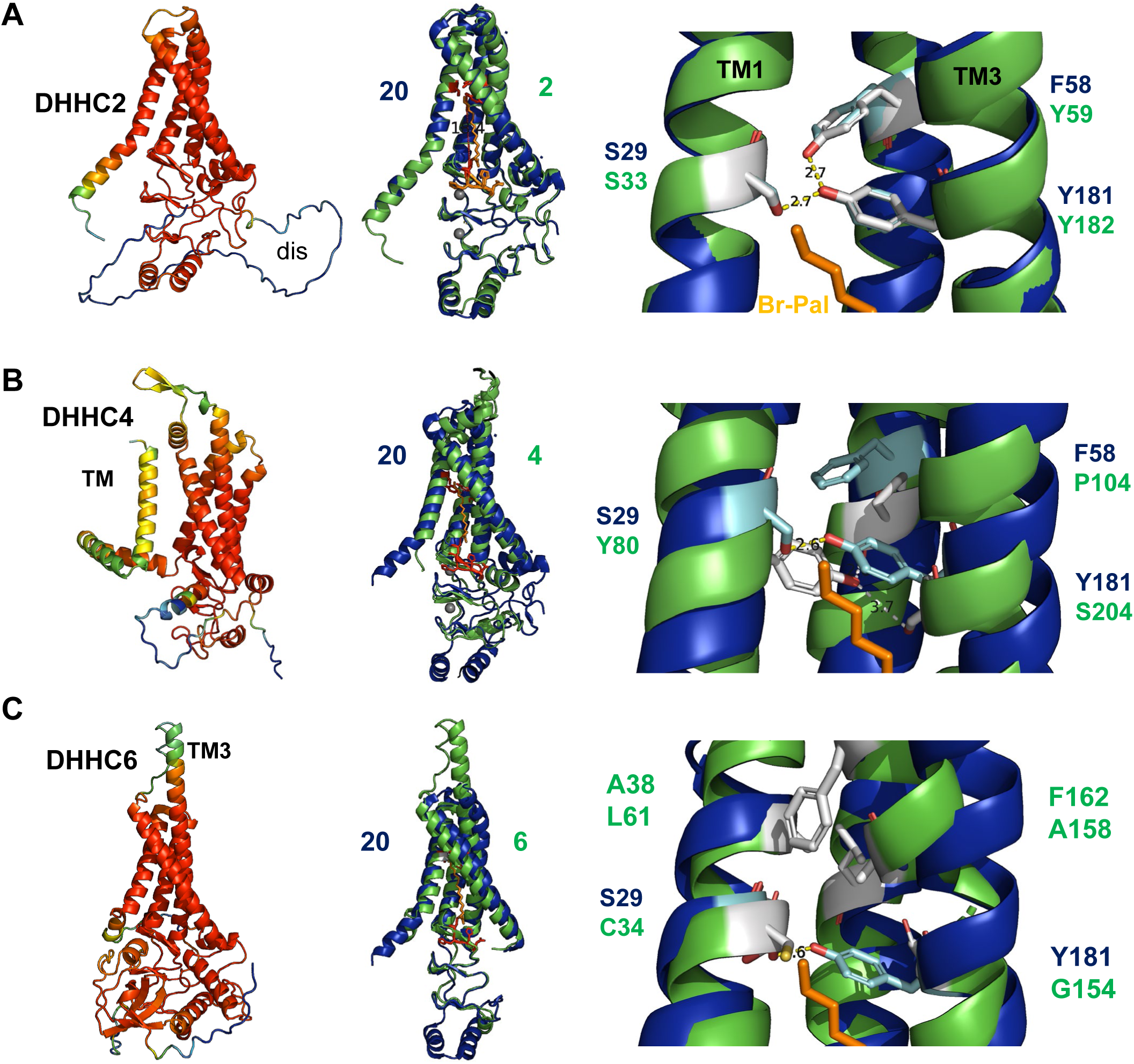
Amino acids at the ceiling of the hydrophobic cavity in DHHC 2, 4 and 6. (A) DHHC 2 structure, (B) DHHC 4 structure, (C) DHHC 6 structure. *Left part*: Alphafold 2 model of the respective human DHHC showing the confidence level of prediction colour-coded from red (high confidence) to blue (low confidence). Dis: disordered region, TM: additional transmembrane region in DHHC 4. TM3: longer transmembrane region in DHHC 6. Accession numbers for the alphafold database: DHHC 2: AF-Q9UIJ5-F1, DHHC 4: AF-Q9NPG8-F1, DHHC 6: AF-Q9H6R6-F1. *Middle part:* Alignment of DHHC 20 with the respective DHHCs. Root-mean-square deviation of atomic positions (RMSD): DHHC 2 = 0.459Å. DHHC 4: 4.326Å, DHHC 6: 2.387Å. The N-terminal extensions in DHHC 2 and DHHC 4 were deleted prior to the alignment. *Right part:* Detail of the alignment showing the amino acids sealing the celling of the cavity as cyan sticks (DHHC 20) or white sticks (DHHC 2, 4 or 6). TM4 was removed from the structure for clarity.

## Discussion

### The function of acylation of Influenza B virus HA

We demonstrate here that palmitate attachment to two conserved cysteine residues in the cytoplasmic tail of HA is essential for Influenza B virus replication. Removing both acylation sites simultaneously or only the C-terminal one prevented recue of infectious virus. Viruses with an exchange of the membrane-near cysteine could be generated but they rapidly reverted back to wild type. The failure to rescue virus particles is not due to instability or reduced transport of HA to the budding site, since removal of both acylation sites does not diminish the amount of HA expressed on the cell surface (Fig. 1, 2). HA from Flu A and B and HEF from Flu C behave identically in this respect, as removal of acylation sites does not affect their plasma membrane transport. [26, 34]. In contrast, acylation of the functionally similar spike protein of SARS-CoV-2 at ten cytoplasmic cysteines is important for the proteińs stability [29].

We have not looked into the particular role of acylation of HA during the viral replication cycle, but in expression experiments deleting one acylation site had an effect on the opening of a fusion pore, which corresponds to a late phase in virus entry [27]. This finding is similar to what has been reported for various subtypes of HA of Flu A and HEF of Flu C virus, although the effect was minor for the latter [18, 21, 22, 26].

The effect of acylation site removal on viral replication distinguishes Influenza A and B viruses from Influenza C viruses. S-acylation of HA of Influenza A and B virus is essential for virus replication since Influenza virus mutants could not be generated if two or three palmitoylation sites were removed. Similarly, rapid reversion of single mutations to cysteines were observed for both viruses. Mutants with one deleted acylation site in the cytoplasmic tail revealed greatly reduced growth rates in cell culture. In contrast, removing the stearoylated cysteine at the end of the TMR of HA (which is not present in HA of Flu B) and also of HEF had only a minor effect [16-18, 24, 26]. Hence, cytoplasmic cysteines connected to palmitate are more critical for virus propagation than transmembrane cysteines bound to stearate. Whether this is due to the specific fatty acid or to the localization of the acylated cysteine is unclear, and also difficult to test experimentally, as shifting a transmembrane cysteine into the cytoplasm also leads to more palmitate being attached [52].

We also found a difference at the intracellular site along the exocytic pathway where HA of Flu A and B are acylated (Fig. 3). Blocking ER exit had no impact on acylation of Flu B HA but lowered Flu A HA acylation to 40%. Thus, acylation of HA of Flu A begins in the ER and proceeds to completion in the Golgi, as has been reported for acylation of the SARS-CoV-2 spike [29]. Both viral glycoproteins are palmitoylated by the enzyme DHHC 20, which is found in both the ER and the Golgi. In contrast, acylation of Flu B HA occurs in the ER and thus corresponds to acylation of the Gp5/M membrane protein of an Arterivirus [53]. The previous findings on the acylation of the hemagglutinating glycoproteins of Influenza viruses A, B and C are summarized in the Fig S4.

### Identification of DHHC enzymes involved in palmitoylation of Influenza B virus HA

The localization of the intracellular acylation site of HA allowed us to narrow down the number of DHHCs possibly involved in palmitoylation. In order to find the enzymes, we have chosen HAP1 cells with individual DHHCs deleted as the experimental system because it yields more consistent results than siRNA-based DHHC knockdown [32]. We started with a screen using cells lacking ER-and Golgi-localized DHHCs. Since the acyl-Rac assay can produce somewhat variable results, we repeated the experiments until the ΔDHHC cells that caused a significant reduction in HA acylation were confidently identified (Fig. 4, 5). Only cells lacking DHHC 1, 2, or 4 showed a decrease in acylation of HA in all six experiments. However, the effect in ΔDHHC 1 cells was small, a reduction of only 10%. In five of six experiments, we observed a substantial decrease in acylation of HA in ΔDHHC 6 cells. However, since in one experiment an increase in acylation was observed, the evaluation of all experiments revealed that the average decrease in acylation was not statistically significantly different from the results with the wt cells. When the first experiment was removed from the analysis, there was a statistically significant difference, and the average reduction in acylation was at the same level as for ΔDHHC 2 and ΔDHHC 4 cells, around 40%. Interestingly, DHHC 6 requires palmitoylation at three cytoplasmic cysteines by DHHC 16 and selenoprotein K as a cofactor to be active [54, 55], although the latter claim is controversial [56] and these dependencies might have caused atypical results.

DHHC 4 and 6 are typical ER-localized proteins with a C-terminal lysine-based retrieval signal and DHHC 1 also resides in the ER by an unknown mechanism [43, 44]. DHHC 2 has been identified in various compartments in different cell types [45, 46]. We show here that all four DHHCs co-localize with HA of Flu B virus and also with an ER-marker in transfected BHK cells (Fig S1, Fig S2). More importantly, DHHC 2, 4 and 6 also resides in the ER in human respiratory A549 cells (Fig. 6), which is consistent with the intracellular organelle where palmitoylation occurs. Since DHHC 4 and 6 are more highly expressed than DHHC 1 and 2 in most human lung cells (S1 table), we assume that they are responsible for the majority of Flu B HA acylation reactions during an Influenza B virus infection. Substrates identified for DHHC 6 include calnexin, CLIMP63 and inositol 1,4,5-triphosphate receptor, which are ER-localized proteins involved in protein folding, ER-morphology and calcium signaling [55, 57, 58]. DHHC 4 palmitoylates proteins of the plasma membrane, such as CD36 and D2 dopamine receptor [59, 60]. DHHC 1 plays a role in STING-dependent innate immune signaling [44] and palmitoylates Interferon-induced transmembrane proteins (IFITM) 1 and 3 [61]. DHHC 2 palmitoylates the tetraspanins CD9 and CD151 [62], members of the src-family kinases [46], and the non-structural protein nsP1 of Chikungunya virus, [63]. Each of these DHHC enzymes thus acylates a variety of proteins with different functions and membrane topologies. However, the exact molecular mechanisms by which the DHHCs recognize their substrate proteins are still largely unknown [3, 4].

### Possible molecular determinants for acylation of Flu A and B HA by different DHHCs enzymes

Although the HA proteins of influenza A and B viruses have similar structure and function, their TMR and short cytoplasmic tail may have minor structural differences that explain acylation by different DHHC enzymes. We predicted the structure of the TMR and cytoplasmic tail of HAs of Flu A and B using alphafold2. Overall, the accuracy of the prediction was quite low, especially for the cytoplasmic tails, which hence might be intrinsically disordered regions. In addition, mutations in the cytoplasmic tail of HA of Flu A and B had no impact on acylation [17, 28, 64]. The middle of the trimeric TMR was predicted with higher confidence, and both the TMRs of Flu A and B contain a region with Ser and Thr residues, flanked by hydrophobic and aromatic residues. However, the backbone of the helices looks different, kinked by the presence of a Gly in the case of the HA of Flu A and straight in the case of the HA of Flu B (Fig S3). Cryo-EM revealed that the transmembrane region of group 1 HA of Flu A splayed out following the Gly [48]. Therefore, the conformation of the TMR, straight or kinked, might allow specific interactions with different DHHCs and present the acylation sites differently to the catalytic domain of DHHCs. The exact mechanism of how DHHC enzymes recognize their substrates is still unknown, but studying the differences between the HAs of Flu A and B may provide insight into this process.

### The specific acylation of HA of Flu B with palmitate is not encoded in the responsible DHHCs

HA of Influenza B virus is known to be exclusively acylated with palmitic acid, unlike HA of Influenza A virus which is acylated with both palmitic and stearic acid [13, 14]. We explored the possible molecular determinants for this fatty acid specificity of HA acylation. One possibility is that DHHC enzymes responsible for attaching the fatty acid to HA may not be able to take up longer-chain fatty acids due to a narrow hydrophobic cavity. However, we found that DHHC 2, 4, and 6 have identical or similar amino acids in the critical position at the end of the transmembrane region as DHHC 20, which can accept both palmitic and stearic acid (Fig. 7). There is also experimental evidence that DHHC 2 and DHHC 4 exhibit no marked fatty acid preference and DHHC 6 can attach stearic acid to one of its substrates [65-67]. Therefore, it is unlikely that DHHC enzymes determine the specificity of HA acylation.

Another possible mechanism is the regulation of the availability of acyl-CoAs for DHHCs. Acyl-CoAs are provided by long-chain acyl-CoA synthetases (ACSLs), transmembrane proteins, of which there are five isoforms in humans. The distinct subcellular localization of different ACSL isoforms could cause the compartmentalization of acyl-CoA [68]. A similar role could be played by acyl-CoA binding domain containing proteins (ACBDs), a large multigene family of diverse proteins. Experimental evidence for the regulation of fatty acid specificity has been provided for ACBD6, which binds to the enzyme N-myristoyltransferase (NMT). The interaction stimulates the transfer of the rare fatty acid myristate, since it locally sequesters Pal-CoA. This explains how the NMT reaction can proceed in the presence of the very abundant substrate Pal-CoA. [69, 70]. Therefore, the connection between protein acylation and lipid metabolism may be stronger than previously thought and requires further investigation [71].

Our study demonstrates the essential role of palmitoylation in the replication of Influenza B virus, which has not been previously studied. Additionally, we identified the responsible DHHC enzymes, which differ from the ones identified previously for acylation of HA of Influenza A virus. Future research can build on these observations by elucidating the specific interactions between viral glycoproteins and DHHC acyltransferases to develop inhibitors that block viral replication.

## Materials and methods

### Cell lines, viruses, genes and plasmids

Human embryonic kidney 293T cells (ATCC CRL-11268), A549 cells (ATCC CCL-185), BHK 21 cells, (ATCC CCL-10), Madin-Darby canine kidney (MDCKII CRL-2936) and HAP-1 cells (Horizon Discovery) were grown in Dulbecco’s modified Eagle’s medium (DMEM; PAN Biotech) supplemented with 10 % fetal bovine serum (FBS), 100 U/ml penicillin and 100µg/ml streptomycin (PS) at 37°C with 5 % CO2. The HAP-1 cells, parental line and knockout cell lines where individual DHHCs were knocked out with CRISPR-Cas9 technology are from Horizon Discovery Bioscience (Vienna, Austria). Editing has been confirmed by Sanger sequencing of genomic DNA. HAP1 is a human near-haploid cell line derived from the chronic myelogenous leukemia (CML) cell line KBM-7. Human Influenza A virus strain A/WSN/33 (H1N1) (Taxonomy ID: 382835) and human Influenza B virus strain B/Lee/40 (Taxonomy ID: 518987) were used in this paper. The 8 pHW2000 plasmids to rescue the Influenza B virus (B/Lee/40 strain), obtained from Thorsten Wolff (Robert Koch-Institute, Berlin), have been described [72]. Mutations in the HA-encoding gene were generated using Quickchange. HA wt and HA Ac1+2 were cloned into the pCAGGS expression plasmid using NheI and XbaI. Mouse DHHC1,2,3,4 and 6 fused at the C-terminus to a HA-tag were kindly provided by Masaki Fukata [73].

### Antibodies and reagents

#### Primary antibodies

Polyclonal antibody from rabbit against H1N1 HA from Influenza A virus WSN (Genetex, GTX127357). Monoclonal antibodies against HA of Influenza B virus were kindly provided by Dr. Florian Krammer (Mount Sinai School of Medicine, New York) [74]. Purified polyclonal Anti-Flotillin-2 from mice (BD, 610383). Polyclonal antibodies to HA tag-ChIP Grade from rabbit (Abcam, Ab9110)

#### Secondary antibodies

anti-mouse IgG (H + L) from goat coupled to horseradish peroxidase, (Biorad, 1706516), anti-rabbit IgG VHH single domain antibody (HRP) coupled to horseradish peroxidase (Abcam, Ab191866). Goat anti-Rabbit IgG (H+L) coupled to Alexa 488 (Invitrogen, # A-11008) Goat anti-Rabbit IgG (H+L) coupled to Alexa 568 (Invitrogen, # A-11011), Goat anti-mouse IgG(H+L) coupled to Alexa 488(Invitrogen, # A28175).

#### Reagents

Lipo3000 transfection reagent (Thermo Fisher, L3000015); Opti-MEM medium (Thermo Fisher Scientific, 31985070); TPCK-trypsin (sigma, T1426); NP40 lysis buffer (Thermo Scientific, 85124); Brefeldin A (Sigma, B7651); Endo H (NEB, P0703L); Pierce ECLplus reagent (Thermofisher, # 32132); SuperSignal West Dura extended Duration substrate (Thermo Fisher #34075); innuPREP Virus TS RNA Kit (Analytik Jena, 845-KS-4710250); High capacity cDNA reverse transcription kit (Thermo Fisher Scientific, #4368814); ER staining kit–Red Fluorescence | Cytopainter (Abcam, ab139482); S7 Fusion Polymerase (Mobidiag, MD-S7-500); Thiopropyl Agarose (Creative biomart, Thio-001A); ProLong Glass antifade mounting medium (Thermo Fisher, P36984); Tris (2-carboxyethyl) phosphine (TCEP, Carl Roth, HN95.2); Methanethiosulfonate (MMTS, Sigma, 208795); Protease inhibitors (Roche/Merck,11873580001) were used in all the protein experiment.

### Transfection-mediated recovery of recombinant Influenza B virus

To generate recombinant Influenza B virus, the eight plasmids pHW-PB2, pHW-PB1, pHW-PA, pHW-HA, pHW-NP, pHW-Lee-NA, pHW-Lee-M, and pHW-Lee-NS (0.5 µg each) were transfected into a co-culture of 293T and MDCKII cells grown in 6-well plates using Lipofectamine 3000. After 5h, Opti-MEM medium was replaced by DMEM medium with 0.1% FBS and 0.5 µg/ml TPCK-trypsin and cells were incubated at 37°C. 72 h after transfection, the supernatant was collected, cell debris were removed by centrifugation (5000rpm, 10min) and the supernatant was kept frozen at -80°C before further usage. One aliquot (500µl, 25%) of this passage 1 (P1) virus was used to infect MDCKII cells in 6 well plates. After 1h the inoculum was removed, cells were washed twice with PBS before adding 2ml infection medium (DMEM with 0.1%FBS and 2 µg/mL TPCK-trypsin). The supernatant was collected after 48h. Another aliquot (200µl) of the P1 virus was injected into the allantois cavity of 9-day-old embryonated chicken eggs which were incubated at 33 °C for 72 hours before the allantois fluid was collected. The resulting P2 viruses from either eggs or MDCK II cells were further passaged using first T25 flasks (P3 virus) and then T75 flasks (P4 virus). TCID50 and HA-assay was used to quantify virus particles.

### RNA extraction, reverse transcription and sequencing of viral HA sequence

200ul cell culture supernatant or allantois fluid was used to extra the RNA using innuPREP Virus TS RNA Kit according to protocol 1 of the manufacturer’s instructions. Reverse transcription was done with the High capacity cDNA reverse transcription kit with RNase Inhibitor (Thermo Fisher Scientific). The reactions were prepared following the manufacturer’s instructions to a total volume of 20µl.The following primers were used to amplify a part of the HA-encoding sequence of Flu B virus. BHA-F: CCAACGAAGGGATAATAAACAGTG, BHA-R: TAACGTTTCTTTGTAATGGTGACAAGC. The S7 Fusion polymerase was used in the PCR reaction and the reaction system was prepared according to the manufacturer’s instructions. The PCR starts at 98°C for 10min and the following program was used for 35 cycles: 98°C for 30s, 55°C for 30 s, and 72°C for 30s, followed by one cycle at 72°C for 10 min. Later the amplified segments were sequenced and compared to the sequence of the wild type.

### Expression of HA and surface trypsinization experiment

70%-80% confluent 293T cells in a 6 well plate were transfected with 3µg HA wt or HA Ac1+2 cloned into the pCAGGS plasmid using lipo3000 transfection reagent. After 48h, the medium was removed, the cells were gently washed twice with PBS and 1ml DMEM without or with 15µg/ml TPCK-trypsin was added to cover the cells. Cells were then incubated at 37℃ for 30min, scraped from the plate, pelleted at 5000rpm for 5min, washed twice with PBS and lysed in 150µl NP40 lysis buffer (final NP40 concentration is 1%) with protease inhibitor. Samples were then analyzed by SDS-PAGE and western blotting

### Brefeldin A experiment

MDCKII cells grown to 90% confluency in 6-well plates (Endo-H experiment) or in T25 flasks (acyl-RAC experiment) were infected with Influenza A or B virus at a MOI of 1. After 1h, the inoculum was removed and cells were washed twice with PBS and 1 ml DMEM with 2% FBS was added. After 2h, Brefeldin A was added to the medium from the stock solution (5 mg/ml) prepared in methanol to the final concentration of 3 µg/ml. Influenza A virus-infected cells were scraped from the plate after 12h and Influenza B virus-infected cells after 24h. Cells were pelleted at 5000rpm for 5min, washed twice with PBS and lysed with 200ul NP40 lysis buffer (final NP40 concentration is 1%) with protein inhibitor on ice for 1h. Insoluble material was pelleted at 10000rpm for 10min at 4°C. Aliquots of the samples were not digested or digested for 3h at 37°C with Endo H according to the manufacturer’s instructions. For the acyl-RAC experiment, the sample from each T25 was lysed on ice for 2h with 1mL buffer A. 10µl 4XSDS-PAGE reducing loading buffer was added and samples heated at 95℃ for 10min prior to western blotting.

### Infection of HAP-1 knockout cells

HAP-1 cells grown in T75 flasks to a confluency of 80% were infected with Influenza B virus at a MOI of 0.5. After 1h, the inoculum was removed, cells were washed twice with PBS and change to 10ml DMDM medium with 2% FBS. After 24h, the cells were collected and centrifugated for 10 min at 5000 rpm to remove the supernatant. The sample from each T75 flask was lysed on ice for 2h with 2ml buffer A. The S-acylation level was measured using Acyl-RAC.

### Acyl-Resin assisted capture (Acyl-RAC) to investigate S-acylation

Protein S-acylation was analyzed by the Acyl-RAC assay as described [75]. Cells were scraped from the plate and lysed in buffer A (0.5%Triton-X100, 25 mM HEPES (pH 7.4), 25 mM NaCl, 1 mM EDTA, and protease inhibitor cocktail) on ice for 2. The cells in T25 flasks were lysed in 1ml buffer A and cells in T75 were lysed in 2ml buffer A. Disulfide bonds were reduced by adding Tris (2-carboxyethyl) phosphine (TCEP) to a final concentration of 20mM and incubated at room temperature for 1h. Free SH-groups were blocked by adding methyl methanethiosulfonate (MMTS, dissolved in 100 mM HEPES, 1 mM EDTA, 87.5mM SDS) to a final concentration of 1.5% (v/v) and incubated for 4 h at 40 °C. 3 volumes of ice-cold 100% acetone were added to the cell lysate and incubated at -20°C overnight. Precipitated proteins were pelleted at 5,000xg for 15 minutes at 4°C. Pelleted proteins were washed five times with 70% (v/v) acetone, air-dried, and then re-suspended in 1ml binding buffer (100 mM HEPES, 1 mM EDTA, 35mM SDS) with protease inhibitor. A small aliquot served as input. Another aliquot of the sample was treated with hydroxylamine (0.5 M final concentration, added from a 2M hydroxylamine stock adjusted to pH 7.4) to cleave thioester bonds. The same volume of the sample was treated with 0.5M Tris-HCl (pH 7.4). 40µl Thiopropyl Agarose, which were washed twice with binding buffer in 1000rpm for 10min before use was added to capture free SH-groups. Samples were incubated with beads overnight at room temperature on a rotating wheel. The beads were then washed 4 times at 5000rpm 5min with binding buffer and proteins were eluted from the beads with 2x reducing SDS-PAGE sample buffer and subsequently heated for 10 minutes at 95°C. Samples were centrifuged at 5000 rpm for 5min. The supernatant was used to detect the S-acylation level by Western blot.

### SDS-PAGE and Western blot

After sodium dodecyl sulfate-polyacrylamide gel electrophoresis (SDS-PAGE) using 12% polyacrylamide, wet electroblotting (100V for 1h) was used to transfer proteins onto polyvinylidene difluoride (PVDF) membranes (GE Healthcare). After blocking (blocking solution: 5% skim milk powder in PBS with 0.1% Tween-20 (PBST)) for 1h at room temperature, membranes were incubated with the primary antibody overnight at 4°C. After washing 3 times for 10min with PBST, membranes were incubated with horseradish peroxidase-coupled secondary antibody for one hour. After again washing three times, signals were detected by chemiluminescence using either the Pierce ECLplus reagent or for weak signals SuperSignal West Dura extended Duration substrate with the Fusion SL camera system (Peqlab, Erlangen, Germany). The density of bands was analyzed with Image J software. Quantification and statistics are fully described in the figure legends.

The first antibodies were used at the following dilutions: monoclonal anti-HA of Flu B from mice: 1:1000, polyclonal antiserum against HA from Influenza A virus H1N1 from rabbit: 1:3000, Purified Anti-Flotillin-2 from mice: 1:1000.

The horseradish peroxidase-conjugated secondary antibodies were used at the following dilutions: recombinant anti-rabbit IgG VHH Single Domain: 1:3000 -1:5000, anti-mouse IgG (H + L) from goat: 1:2000 -1:3000. The primary and secondary antibodies were diluted in PBS containing 3% BSA and 0.1% tween.

### Fluorescence microscopy of transfected and infected cells to generate recombinant Flu B virus

48h post-transfection or infection, cells were fixed with 4% paraformaldehyde (PFA) for 20min at room temperature, washed two times with PBS, permeabilized with Triton X-100 (0.1% in PBS) for 10 min and again washed twice with PBS. Cells were stained with mouse monoclonal antibody against Influenza B virus HA at a dilution of 1:1000 for 1h. Cells were washed 3 times for 5 min with PBS and later covered with anti-mouse coupled to alexa 488 fluorescent antibody (1:1000) for 1h. Cells were washed 3 times for 5 min with PBS. Cells were illuminated via laser lines at 488 nm (Alex Fluor 488) and visualized with ZEISS Colibri7, 20X objective was used.

### Confocal microscopy to co-localize DHHCs with the endoplasmic reticulum or with HA of Flu B

#### Colocalization with the endoplasmic reticulum

BHK 21 and A549 cells were seeded at 50% confluency one day before transfection on glass coverslips in 24-well cell culture plates. Cells were transfected with 500ng plasmids encoding mouse DHHC 1, 2, 3, 4 or 6 fused at the C-terminus to a HA-tag with using lipofectamine 3000 transfection reagent according to manufacturer instructions. 24h post-transfection, cells were fixed with 4% paraformaldehyde (PFA) for 20min at room temperature, washed twice with PBS, permeabilized with Triton X-100 (0.1% in PBS) for 10 min and again washed twice with PBS. To coat non-specific protein binding sites, cells were incubated with bovine serum albumin (BSA, 3% in PBST) for 1h at room temperature. BSA was removed and cells were incubated with the primary rabbit polyclonal antibody to HA tag (1:500 dilution in 3%BSA) for 1h at room temperature. After washing three times with PBS for 5 min, cells were incubated with anti-rabbit coupled to Alexa 488 fluorescent secondary antibody (1:1000) in the dark. Cells were washed 3 times for 5 min with PBS. The ER was stained with staining kit–Red Fluorescence Cytopainter at 1:1000 dilution (with 1X assay buffer from the kit) for 15 min at 37°C. Cells were washed 3 times for 5 min with PBS

#### Colocalization with HA

BHK 21 cells were seeded at 50% confluency one day before transfection on glass coverslips in 24-well cell culture plates. Cells were co-transfected with 250ng the HA gene in pCAGGS and 250ng plasmids encoding mouse DHHC 1, 2, 4, and 6 fused with HA tag using lipofectamine 3000 transfection reagent according to manufacturer instructions. After 24 h cells were washed and fixed and then incubated with the rabbit polyclonal antibody to the HA-tag (1:500 dilution in 3%BSA) for 1h at room temperature and later with anti-rabbit coupled to Alexa 561 fluorescent secondary antibody (1:1000). HA was detected by the monoclonal antibody against Influenza B virus HA (1:1000) and anti-mouse coupled to alexa 488 fluorescent secondary antibody (1:1000). Cells were subsequently stained with Hoechst 33342 from the kit (1:2000 dilution in 1X assay buffer) for 10 min at room temperature to visualize nuclei. Cells were washed 3 times with PBS and coverslips were mounted on glass slides with ProLong Glass antifade mounting medium (Thermo Fisher) and allowed to cure in a dark place overnight.

Cells were illuminated via laser lines at 405nm (Hoechst 33342), 488 nm (Alex Fluor 488) and 561 nm (ER-staining and Alex Fluor 568) and visualized with the Leica Stellaris 8 FALCON Confocal Microscope (Leica) using the objective: HC PL APO 63x/1.40 OIL SC2. WD 0.14 mm. The images were then processed using Fiji software.

### Predicted structures of DHHCs and of the C-terminus of HA using Alphafold2

The predicted structures of human DHHCs were downloaded from the alphafold protein structure database: https://alphafold.ebi.ac.uk/. The accession numbers are: DHHC 2: AF-Q9UIJ5-F1, DHHC 4: AF-Q9NPG8-F1, DHHC 6: AF-Q9H6R6-F1.

Since no predicted structures of viral proteins are in this database, we used CoLabFold V1.5.2. to make our own prediction of the C-terminus of HA of Flu A and B: (https://colab.research.google.com/github/sokrypton/ColabFold/blob/main/AlphaFold2.ipynb The amino acid sequence used for the predictions are shown in the legend of Fig S1.

Alphafold2 generates a PDB file and a confidence score, called the predicted local distance difference test (pLDDT), which indicates the confidence in the local structure prediction. The scale ranges from 0 to 100 and an IDDT value above 90 indicates very high accuracy, equivalent to structures determined by experiments, which allows for the investigation of details of individual side chains. A value from 70 to 90 indicates high accuracy, where the predictions of the protein’s backbone are reliable. A value of 50 to 70 indicates lower accuracy, but the predictions of the individual secondary structural elements, α-helices and β-strands are probably correct, but how they are aligned in space is uncertain. Values below 50 might be an indication of an intrinsically unstructured region. The PDB files with the predicted structure contain this information in the B-factors, which can be highlighted in the 3D protein structure. Areas with high B-factors, which indicates high confidence, are colored red, while low B-factors are colored blue.

The figures were generated with PyMol (Molecular Graphics System, Version 2.0 Schrödinger, LLC, https://pymol.org/2/).

## Acknowledgments

We thank Thorsten Wolff (Robert-Koch Intitute, Berlin) for the Influenza B virus reverse genetics plasmids, Florian Krammer (Mount Sinai School of Medicine, New York) for monoclonal antibodies against Flu B HA and Masaki Fukata (National Institutes for Physiological Sciences, Japan) for DHHC clones. We thank the Service Unit Microscopy of our faculty for help with confocal microscopy.

M.V. designed and supervised the study and wrote the manuscript. X.M. performed the experiments and revised the manuscript.

This work was supported by the German research foundation [grant no: Ve 141/18-1] to Michael Veit. Xiaorong Meng is the recipient of a Chinese scholar council (CSC) fellowship. The funders had no role in the study design, data collection and analysis, decision to publish or preparation of the manuscript.

## Data availability

All data generated during this study are included in the manuscript and supporting files.

---

**Fig S1.**
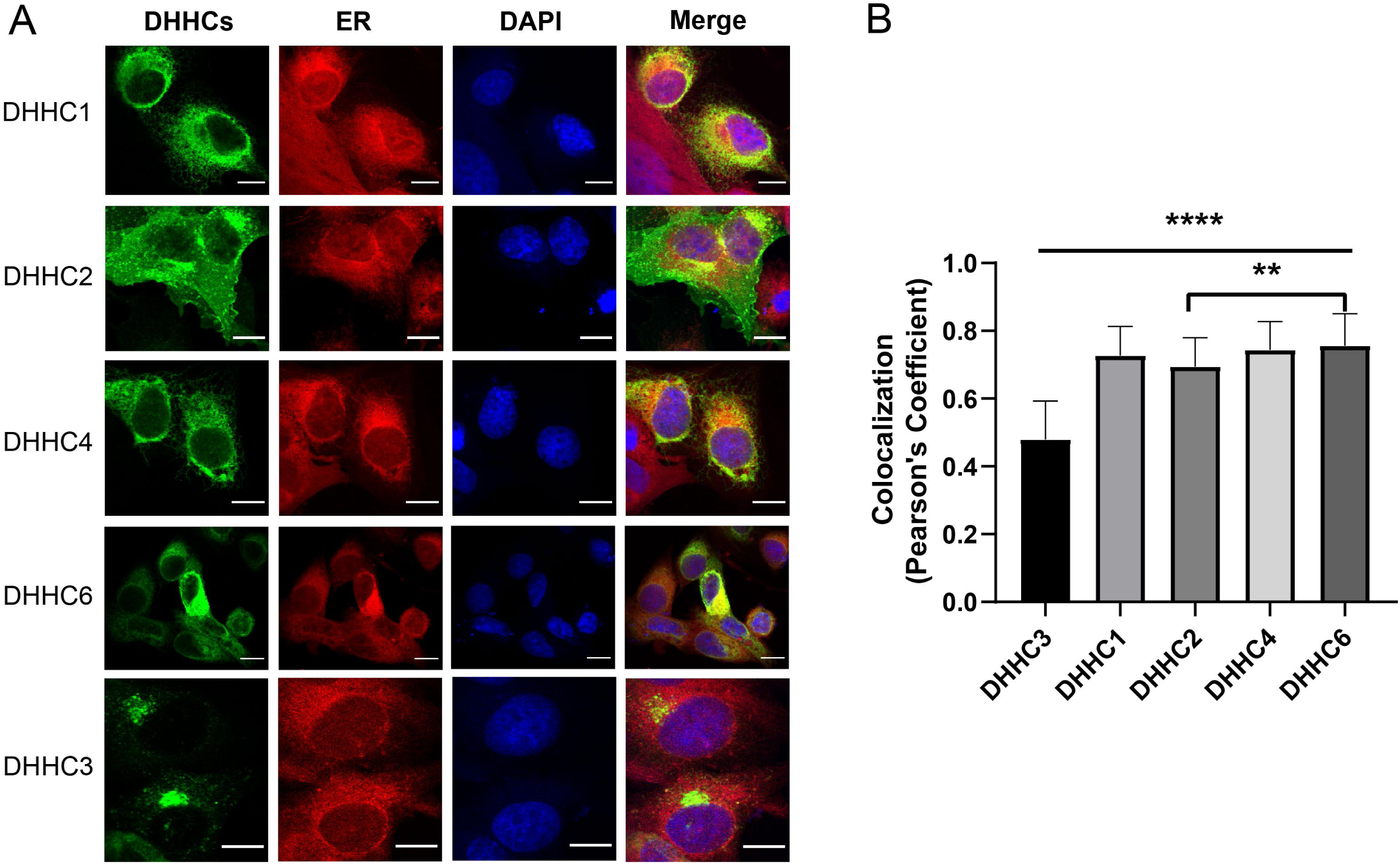
Co-localisation of DHHCs responsible for acylation of HA with ER marker. (A) Confocal microscopy of BHK cells transfected with the indicated mouse DHHCs fused to a HA tag. Cells were permeabilized and stained with antibodies against HA tag (green), with ER staining kit (red), and with DAPI (blue) to highlight the nucleus. The scale bar is 10 µm. (B) Co-localization of DHHCs and ER from at least 50 cells was quantified with Pearson’s correlation coefficient method using the JACoP plugin of the Fiji software. One-way ANOVA followed by multiple comparison Tukey test was applied for statistical analysis. ****, P <0.0001 DHHC 3 versus DHHC 1,2,4 and 6; **, P <0.01 DHHC 2 versus DHHC 6. DHHC 1,4,6 has no significant differences from each other. There is no significant difference between DHHC 1, DHHC 4, and DHHC 6.

**Fig. S2.**
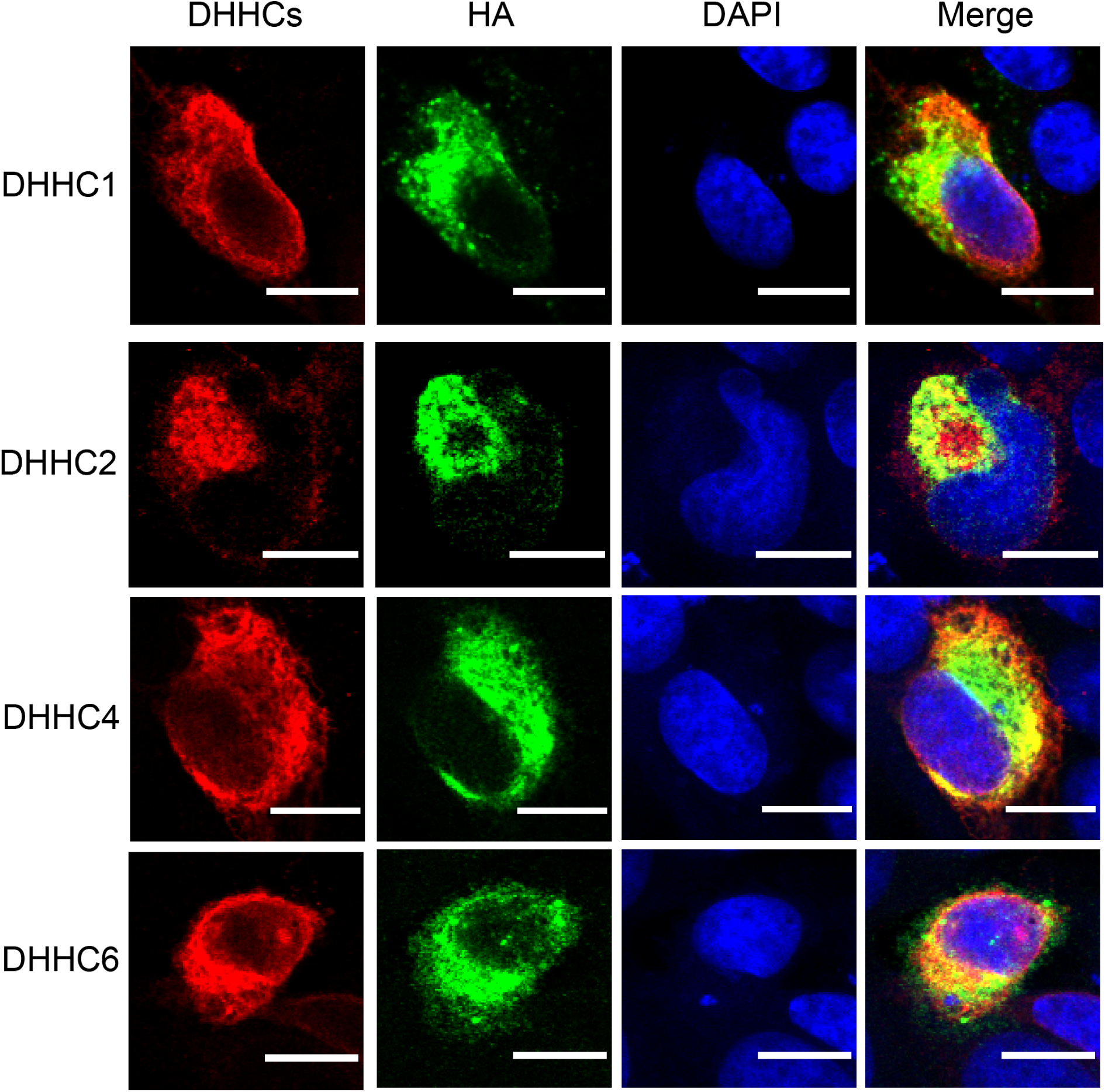
Co-localisation of DHHC 1,2,4 and 6 with Flu B HA. Confocal microscopy of BHK cells co-transfected with the indicated mouse DHHCs fused to a HA tag and Influenza B virus HA. Cells were permeabilized and stained with antibodies against HA-tag (red) and against Flu B HA (green), and with DAPI (blue) to highlight the nucleus. The scale bar is 10 µm.

**Fig. S3.**
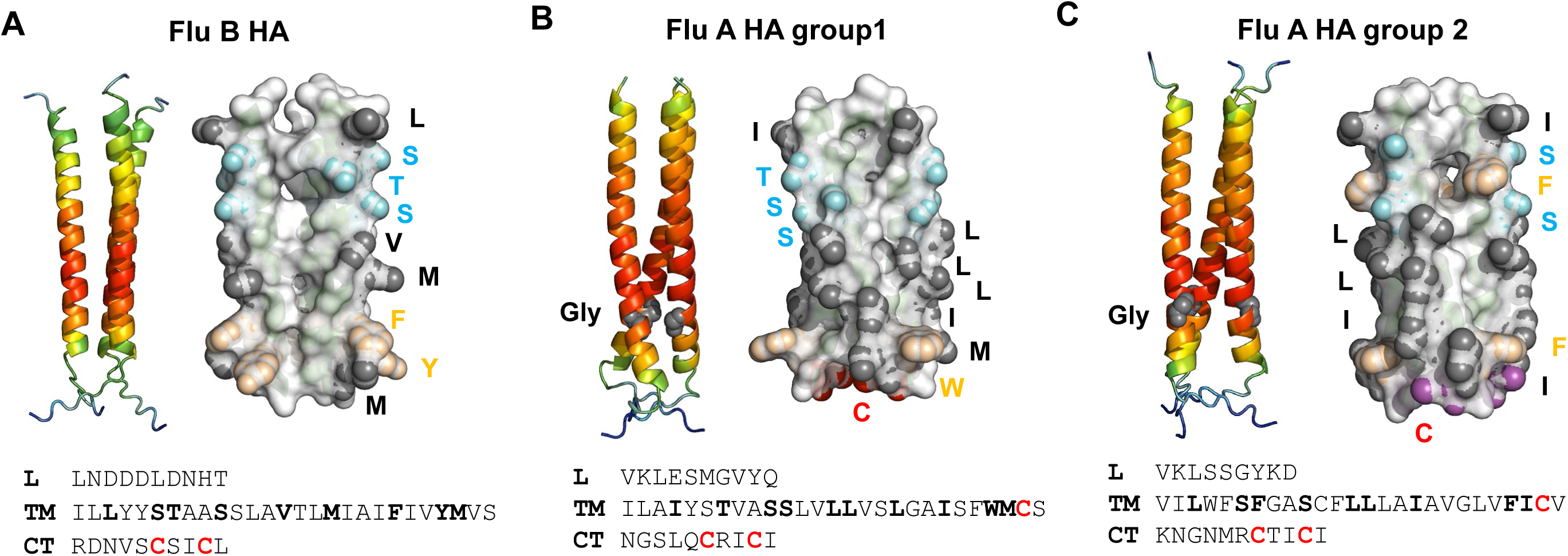
Predicted structure of the transmembrane region of HA of Flu A and B. (A) Flu B HA (B) Flu A HA group 1 (C) Flu A HA group 2. Left part: Cartoon of the structure predicted by alphafold2. The confidence score is colour coded from red (high confidence) to blue (low confidence). Right part: Surface representation of the transmembrane region. Amino acid side chains located at the surface are highlighted as spheres. Cyan spheres: Hydroxy amino acids, black spheres: long and hydrophobic amino acids, wheat spheres: aromatic amino acids. Lower part: amino acid sequence used for the prediction divided into L: linker region, TM: transmembrane, CT: cytoplasmic tail. Amino acids highlighted in the surface representation are bold, the acylated cysteines in red. Sequences are from Flu B Lee strain, Flu A/WSN/1933(H1N1) group 1 HA, Flu A/FPV/Rostock/34, H7N1 group 2 HA.

**Fig S4.**
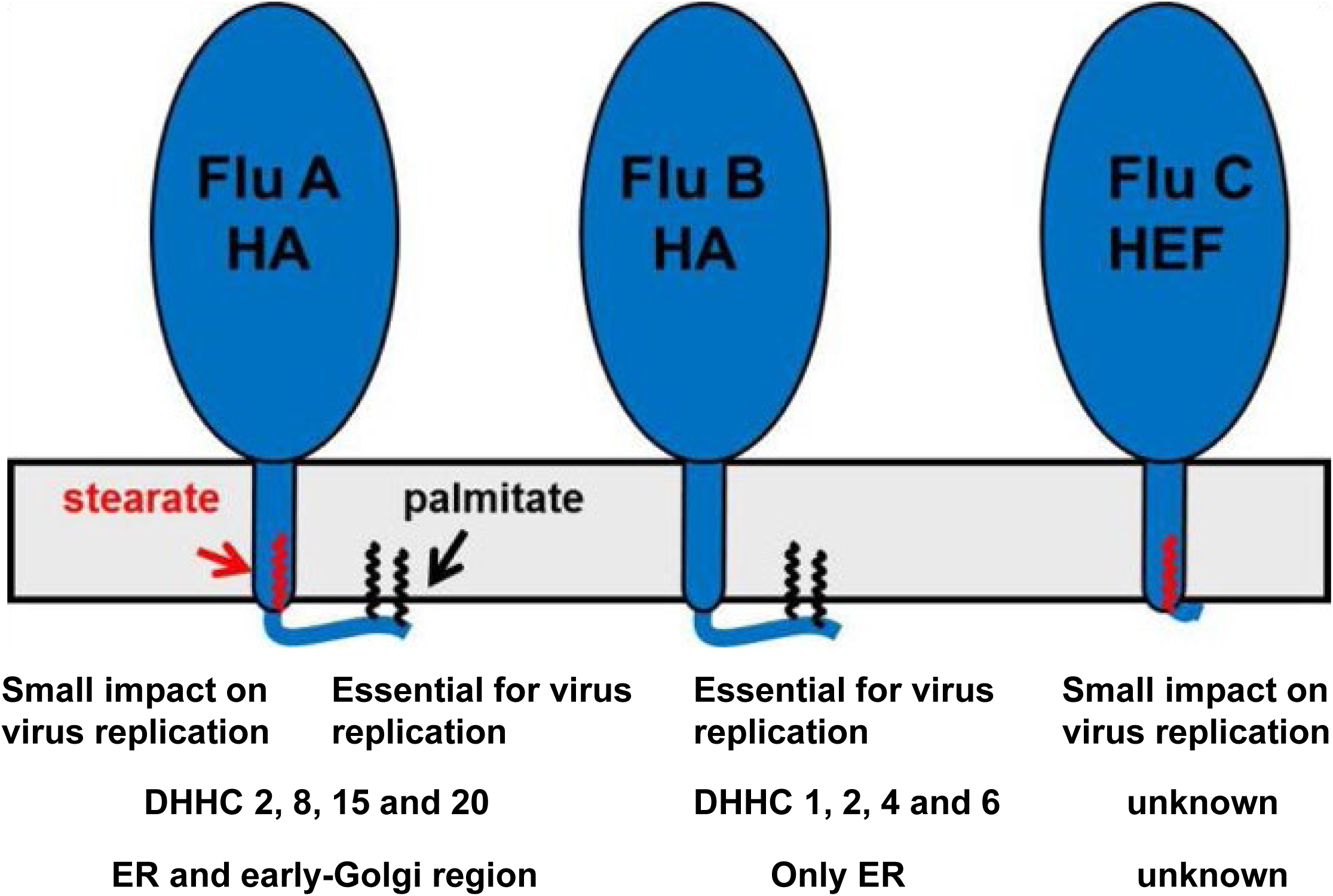
Summary of results on acylation of hemagglutinating glycoproteins of Influenza A, B and C virus. Schematic of hemagglutinating glycoproteins HA and HEF embedded within a membrane. Fatty acids linked to cysteine residues are shown as red (stearate) or blue (palmitate) zigzag line. The impact of removing individual acylation sites, the DHHCs involved in acylation and the intracellular site are listed. In addition, whereas HA of Influenza A and B virus are associated with membrane rafts, cholesterol- and sphingolipid-enriched nanodomains of the plasma membrane, HEF is thought to localize to the bulk phase of the plasma membrane

**Table S1.**
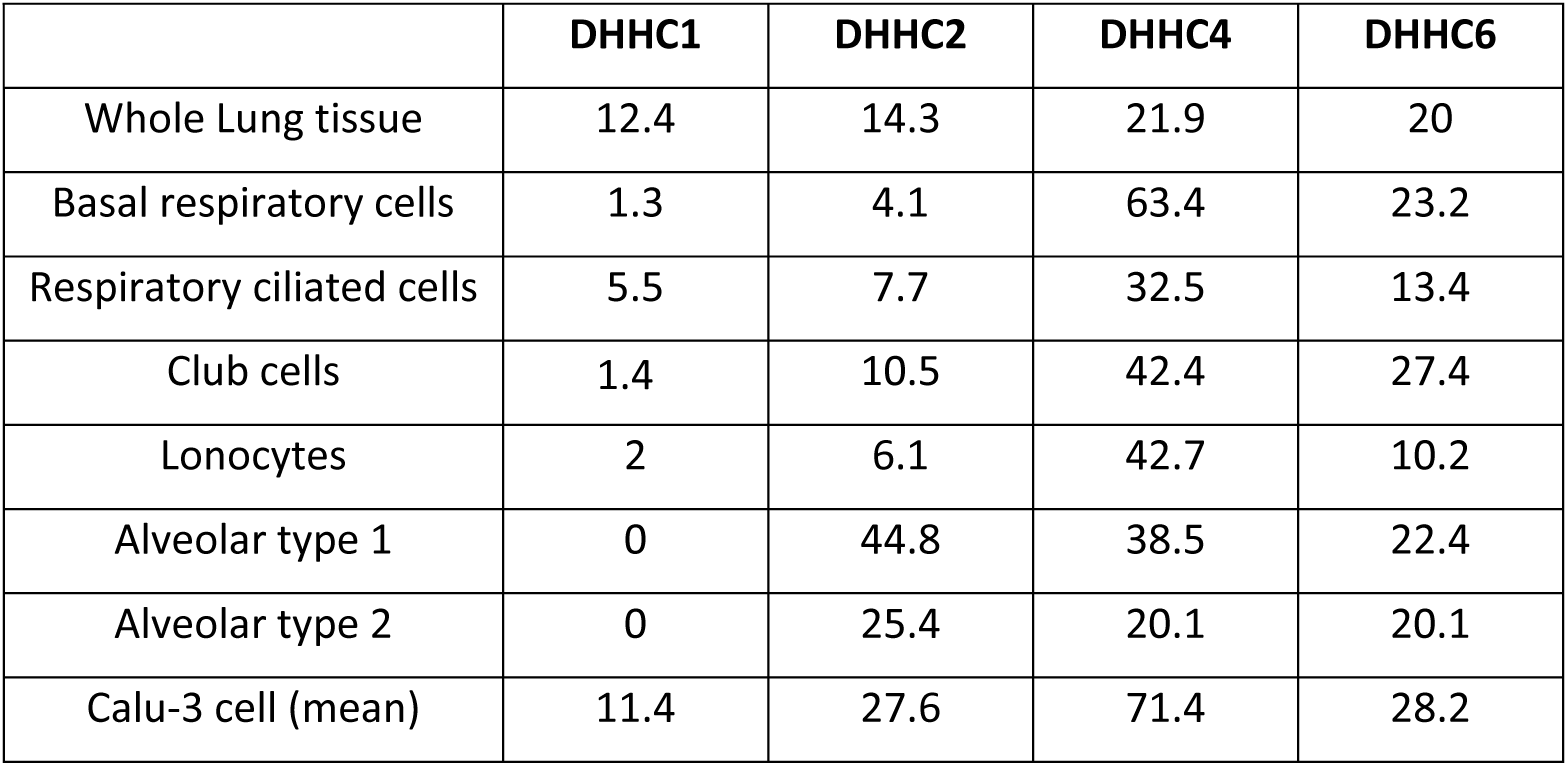
Expression of DHHC 1, 2, 4 and 6 in human lung tissue and various cell types. Expression levels of the indicated DHHCs in whole lung tissue and specialized lung cell types. The results are based on RNA-Seq experiments and the units are transcripts per million (nTPM). Data were taken from the “CellPalmSeq” database (https://cellpalmseq.med.ubc.ca/), which is based on the human protein atlas (https://www.proteinatlas.org/

